# The Rap1 GTPase performs two separable functions during cellularization to enable ventral furrow formation

**DOI:** 10.64898/2026.09.21.753314

**Authors:** Amruta P Nayak, Michael Glotzer

## Abstract

Morphogenetic processes involve dynamic cell-level events, such as contractility, shape changes, and adhesion between cells and extracellular structures. These cell-level changes require intercellular coordination in order to generate the appropriate tissue-level output. Although the repertoire of these cell-level changes is limited, these processes are reused serially during development to generate diverse tissue-level transformations. Dissecting any one of them benefits from perturbations controlled with high spatiotemporal resolution. Despite extensive analysis of ventral furrow formation, a model gastrulation event in the early Drosophila embryo, it remains unclear how tissue-level events are orchestrated at the molecular level. We therefore developed a biosensor and optogenetic tools for spatiotemporal modulation of Rap1, a small GTPase known to be required for ventral furrow formation. Using the biosensor, we show that Rap1 activity declines in ventral cells as the furrow forms. Optogenetic dissection of Rap1 activity reveals that acute inhibition during furrowing has little effect. Instead, Rap1 contributes to furrowing through two separable functions during the preceding process, cellularization: it is continuously required for efficient membrane ingression, and transiently required to nucleate the adherens junctions that later coordinate contractility across the ventral epithelium.

## Introduction

During early embryogenesis, multiple processes occur in parallel or close succession to generate different embryonic features. These processes, including cell division, zygotic gene activation, cell fate specification, and morphogenesis, are governed by distinct regulatory pathways, yet they must be coordinated in space and time to produce a functional organism. This coordination is particularly important in early embryonic development, in which the product of one stage becomes the substrate for the next. In the *Drosophila* embryo, morphogenetic events occur in rapid succession to position a set of cells or specify their fates at the correct location within the embryo. First, cellularization converts the syncytial blastoderm into a continuous epithelium (Schejter and Wieschaus, 1993, Lecuit and Wieschaus, 2000). Ventral furrow formation begins immediately afterward, internalizing the presumptive mesoderm within 15 minutes (Leptin and Grunewald, 1990, Sweeton et al., 1991, Martin et al., 2009). Continuing this rapid, stereotyped sequence, the germ band then extends as mesoderm internalization completes (Foe, 1989, Irvine and Wieschaus, 1994, Bertet et al., 2004, Martin et al., 2009). Each of these processes involves myosin-driven mechanical deformation of the epithelium (Martin et al., 2009, Rauzi et al., 2010). In addition, cells in the epithelium are connected to their neighbors through adherens junctions that are established *de novo* during cellularization and remodeled as the epithelium transforms (Müller and Wieschaus, 1996, Bertet et al., 2004, Rauzi et al., 2010). These junctions are mechanosensitive and linked to the actomyosin cytoskeleton, allowing contractile forces to be integrated over supracellular length scales (Cavey et al., 2008, Martin et al., 2010, Lecuit and Yap, 2015). Junction assembly and remodeling are therefore critical to morphogenesis (Harris and Tepass, 2010).

The small GTPase Rap1 is a conserved regulator of cell adhesion that is known for its role in positioning cadherin-based cell-cell adhesion and integrin-mediated cell-matrix adhesion in diverse contexts (Knox and Brown, 2002, Boettner et al., 2003, Jaśkiewicz et al., 2018). In both settings, it recruits cytoskeletal linker proteins that connect the adhesion complexes to actin: Canoe/Afadin to adherens junctions and Talin to focal adhesions either directly or through Rap1-GTP-interacting adapter molecule (RIAM) (Boettner et al., 2003, Sawyer et al., 2009, Yang et al., 2014, Zhang et al., 2014, Plak et al., 2016, Bromberger et al., 2018). During cellularization of the *Drosophila* embryo, Rap1 and Canoe are coordinately required for apical positioning of Bazooka/Par3 and adherens junctions (Choi et al., 2013). At this stage, Rap1 itself is distributed uniformly along the lateral membrane, leading to the hypothesis that Rap1 activity, rather than its localization, is functionally relevant and spatially regulated (Bonello et al., 2018). Consistent with this, two Rap1 guanine nucleotide exchange factors (GEFs) act during early embryogenesis: the ELMO-Sponge complex is required for the initial subapical restriction of Canoe at the onset of cellularization, and Dizzy regulates Canoe enrichment at junctions during late cellularization (Spahn et al., 2012, Bonello et al., 2018, Schmidt et al., 2018).

Ventral furrow formation is the first morphogenetic movement shown to require Rap1 (Asha et al., 1999). During this process, loss of either Rap1 or its activity (by loss of its GEF Dizzy) produces apical constrictions that are slow and poorly coordinated, resulting in a failure to furrow (Spahn et al., 2012). The defects are not due to a failure in patterning, since ventral (Twist, Snail) and terminal (Huckebein) markers are unaffected by these perturbations (Spahn et al., 2012). Subsequent morphogenetic events, namely posterior midgut invagination and germ-band extension, still commence, indicating that the onset of these later processes is unaffected. The preceding stage, cellularization, also appears largely normal. Actomyosin organization is intact and the cellularization fronts are reported to reach the basal end to form complete cells. The apical adherens junctions, however, are an exception. Upon Rap1 depletion, junction components show a fragmented and reduced apical localization during cellularization and initially fail to form a continuous circumferential belt (Spahn et al., 2012, Choi et al., 2013). By the onset of ventral furrowing, however, this localization is partially restored (Choi et al., 2013). Given that adherens junctions anchor the actomyosin cytoskeleton, disrupted junctions have been proposed to compromise force transmission and produce poorly coordinated constrictions (Sawyer et al., 2009, Martin et al., 2010, Spahn et al., 2012). Although these data establish that Rap1 activity is essential for ventral furrow formation, it is not clear when and where Rap1 is functional.

Our limited understanding of when Rap1 activity is required can be ascribed to three factors. First, Rap1 function has been dissected primarily with chronic perturbations: germline clones, maternal/zygotic null mutants, RNAi, and constitutively active or dominant-negative variants (Asha et al., 1999, Spahn et al., 2012, Choi et al., 2013). None of these allows probing of activity dynamics. Second, the mutants have been largely assayed by fixed imaging at discrete developmental stages, which cannot capture the emergence of the phenotypic changes. This is particularly limiting in early embryonic development, where events occur in rapid succession and may not be fully independent of one another. Third, small GTPases like Rap1 are switch-like molecules. Their function depends not only on abundance but also on regulated cycling between GTP- and GDP-bound (active and inactive) states, which is not preserved in chronic perturbations. Indeed, some studies report similar phenotypes with both constitutively active and dominant-negative mutants, and therefore do not clarify how Rap1 activity affects the process (Wang et al., 2013). Addressing these limitations requires perturbations that are acute and reversible. Additionally, a means of detecting the active form is necessary to establish the profile of Rap1 activity across these stages.

We therefore built a toolkit for acute, reversible control and direct readout of Rap1 activity in the *Drosophila* embryo: optogenetic tools for stage-restricted perturbation, and a localization-based biosensor that reports Rap1 activity in live embryos. Using these tools, we find that Rap1 has two separable functions during cellularization. First, Rap1 acts continuously to maintain the rate of membrane ingression. Second, Rap1 acts during early cellularization to establish apical adherens junctions, a process critical for ventral furrow formation. In contrast to these early requirements, Rap1 activity is largely dispensable during ventral furrowing itself.

## Results

### Maternal Rap1 depletion slows cellularization and delays ventral furrow formation

Prior characterizations of the Rap1 requirement in ventral furrow formation (hereafter VFF) largely relied on fixed imaging of individual stages (Asha et al., 1999, Spahn et al., 2012, Choi et al., 2013). As these processes are highly dynamic, we used live imaging to follow the kinetics of cellularization and VFF, asking whether the reported VFF defect reflects a direct requirement during furrowing or an indirect consequence of an earlier defect. Maternal Rap1 was depleted using RNAi, previously shown to deplete Rap1 in the early embryo (Sheppard et al., 2023).

We imaged embryos co-expressing Jupiter-GFP and Gap43-mCherry, which report on microtubule organization and the cell membrane, respectively (Fig. 1A, B; Videos S1 and S2). We tracked cell-cycle progression as an independent readout of developmental timing by visualizing the organization of the perinuclear ring of microtubules. Through the first hour after anaphase onset of nuclear cycle 13 (NC13), spanning most of cellularization, cell-cycle progression is indistinguishable between control and Rap1-depleted embryos. The perinuclear microtubule rings undergo stereotyped, cell-cycle-linked changes (Warn and Warn, 1986, Foe, 1989), with no difference between genotypes (Fig. 1C). The assay thus detects the previously reported cell-cycle dynamics and shows that they are unaffected by Rap1 depletion.

**Figure 1.**
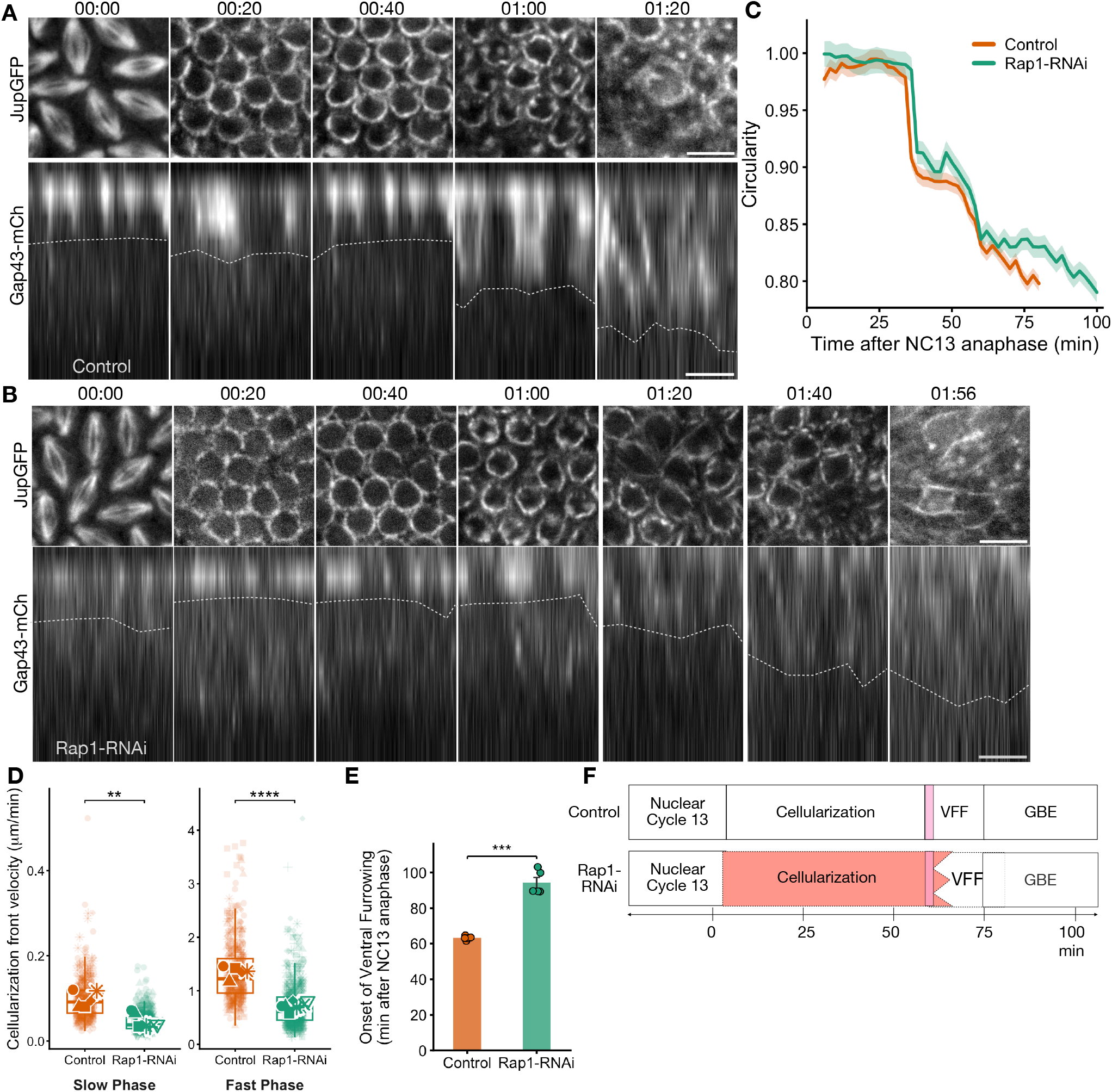
Maternal Rap1 depletion slows cellularization and disrupts ventral furrow formation. (A, B) Time-lapse images of control (A) and maternal Rap1-RNAi (B) embryos, expressing Jupiter-GFP (microtubules; *en face*, top) and Gap43-mCherry (plasma membrane; cross-section, bottom, apical up) across nuclear cycle 13 (NC13), cellularization, and ventral furrow formation (VFF). Time (hh:mm) is relative to anaphase onset of NC13. Dashed gray lines indicate the tips of cellularization fronts. The Rap1-RNAi series extends to 01:56 because cellularization and furrow onset are delayed. Scale bars, 10 μm. (C) Apical microtubule ring organization as measured by circularity over time in control (orange) and Rap1-RNAi (green) embryos. Circularity (4πA/P², where P is ring perimeter and A the enclosed area) equals 1 for a circle and decreases as the ring becomes convoluted. Lines, per-genotype mean; shaded band, SEM across embryos; n = 5 embryos per genotype, median 31 rings per embryo per time point. Time is relative to anaphase onset of NC13. Traces extend to different endpoints because cellularization and ventral furrow formation are delayed in Rap1-RNAi embryos. (D) Cellularization front velocity during the slow and fast phases in control (orange) and Rap1-RNAi (green) embryos. Each point is one measured cellularization front; point shape denotes embryo; corresponding larger shapes with white borders denote per-embryo means; boxes, median and interquartile range (IQR) of pooled measurements; whiskers, 1.5×IQR. Welch’s t-test on per-embryo means. Note the different y-axis scales for the two phases. (E) Onset of ventral furrowing, scored as the interval between anaphase onset of NC13 and the first apical constrictions visible in the ventral epithelium. Bars, mean; error bars, SEM; points, individual embryos. Welch’s t-test, p < 0.001. (F) Schematic of early embryonic developmental events in control and Rap1-RNAi embryos. Pink bar, the end of the interval over which cell-cycle progression is equivalent in the two genotypes (C). Up to this point, development proceeds on the same schedule; beyond it, slowed cellularization delays the onset of apical constriction (E) and therefore of ventral furrow formation (VFF, red), which extends into the window normally occupied by germ-band extension (GBE).

Unlike the cell cycle, membrane ingression is slowed by Rap1 depletion. Ingression during cellularization is normally biphasic, comprising slow and fast phases (Lecuit and Wieschaus, 2000). In controls, cellularization proceeds with morphology and kinetics similar to those previously reported (Merrill et al., 1988, Foe, 1989, Figard et al., 2013, He et al., 2016) (Fig. 1A, bottom). Upon Rap1 depletion, which results in a strong reduction in cortical signal from the Rap1 activity biosensor (see below) (Fig. 1—Supplement 1), ingression is slower overall (Fig. 1B, bottom), with the rates of both phases roughly halved (Fig. 1D). Rap1-depleted embryos also show a substantially longer interval between anaphase onset of NC13 and the first apical constrictions in the ventral epithelium (Fig. 1E). These embryos subsequently fail to complete VFF (Fig. 1—Supplement 2). Rap1 depletion thus slows cellularization without delaying the cell cycle.

Cellularization does eventually reach completion upon Rap1 depletion and VFF onset is delayed relative to anaphase onset of NC13 (Fig. 1F). These temporal shifts result in an additional complication. Because the onset of VFF is delayed, it may overlap with the developmental window normally occupied by germ-band extension. Elongation of the tissues generates posterior-directed forces that could impact completion of ventral furrowing. Moreover, because chronic depletion removes Rap1 globally, throughout development, such experiments do not reveal where and when Rap1 functions. To alleviate these issues, and to more precisely identify the spatiotemporal requirements of Rap1 function, tools capable of spatiotemporal perturbations are required.

### A biosensor and optogenetic probes enable acute measurement and manipulation of Rap1 activity *in vivo*

To directly visualize Rap1 activity, we first adapted an existing mammalian localization-based reporter for use in *Drosophila*. This reporter uses the Ras-association (RA) domain, also called the Ras-binding domain (RBD), from the mammalian Rap1 effector RalGDS. The RA domain is present in numerous Rap1 effectors and has a higher affinity for the GTP-bound vs. GDP-bound Rap1 (Franke et al., 1997). Tagged with GFP, it has been used to report Rap1 activity during live imaging of cultured mammalian cells (Bivona et al., 2004). We generated a transgenic line ubiquitously expressing GFP-RalGDS^RA^. This reporter accumulates at the cortex in the early embryonic epithelium. When Rap1 is depleted or inactivated (maternal Rap1 RNAi or dominant-negative Rap1^S17A^ overexpression, respectively; Fig. 2—Supplement 1A), cortical accumulation of GFP-RalGDS^RA^ is reduced but not eliminated (see inset), though little Rap1-dependent signal should remain. The residual signal compresses the reporter’s dynamic range, making modest changes in Rap1 activity difficult to resolve.

We therefore tested whether the RA domain of *Drosophila* Rgl would function as a better reporter in this system. Rgl is the *Drosophila* homolog of mammalian RalGDS. Their RA domains share substantial sequence identity (Mirey et al., 2003) and the predicted Rgl^RA^ domain superposes closely on the experimentally determined RALGDS domain (Fig. 2— Supplement 1D). We tagged this domain with dTomato, chosen for compatibility with the blue-light-activated optogenetic tools described below, and expressed it ubiquitously (*Ubi>*dTom-Rgl^RA^; hereafter Rap1-biosensor). Homozygous Rap1-biosensor flies are viable and fertile, indicating that ubiquitous expression is well tolerated. The Rap1-biosensor localizes to the cortex in the embryonic epithelium, the expected site of Rap1 activity, as Rap1 is geranylgeranylated and membrane-associated (Kawata et al., 1990) (Fig. 2A, 2B left). To test whether cortical biosensor localization detects active Rap1, we altered maternal Rap1 levels using RNAi or impaired Rap1 cycling by overexpressing nucleotide-locked forms, and measured cortical biosensor enrichment relative to the Rap1-independent membrane marker Spider-GFP. Reducing Rap1 levels by RNAi decreases the cortical biosensor enrichment by nearly half. Overexpressing dominant-negative Rap1^S17A^ similarly reduces cortical biosensor enrichment, though to a lesser extent. Conversely, overexpression of constitutively active Rap1^Q63E^ enhances cortical association of the sensor. Because the measured changes in biosensor signal are normalized to Spider-GFP, they are independent of changes in membrane morphology. The bidirectional response therefore indicates that cortical recruitment of the biosensor is sensitive to Rap1 activity (Fig. 2B, 2C). Under similar conditions, dTom-Rgl^RA^ outperforms GFP-RalGDS^RA^: it shows a larger decrease in cortical enrichment upon decreased Rap1 levels or activity and lower residual signal, indicating improved dynamic range and less Rap1-independent background in the early embryo (Fig. 2—Supplement 1A-C).

**Figure 2.**
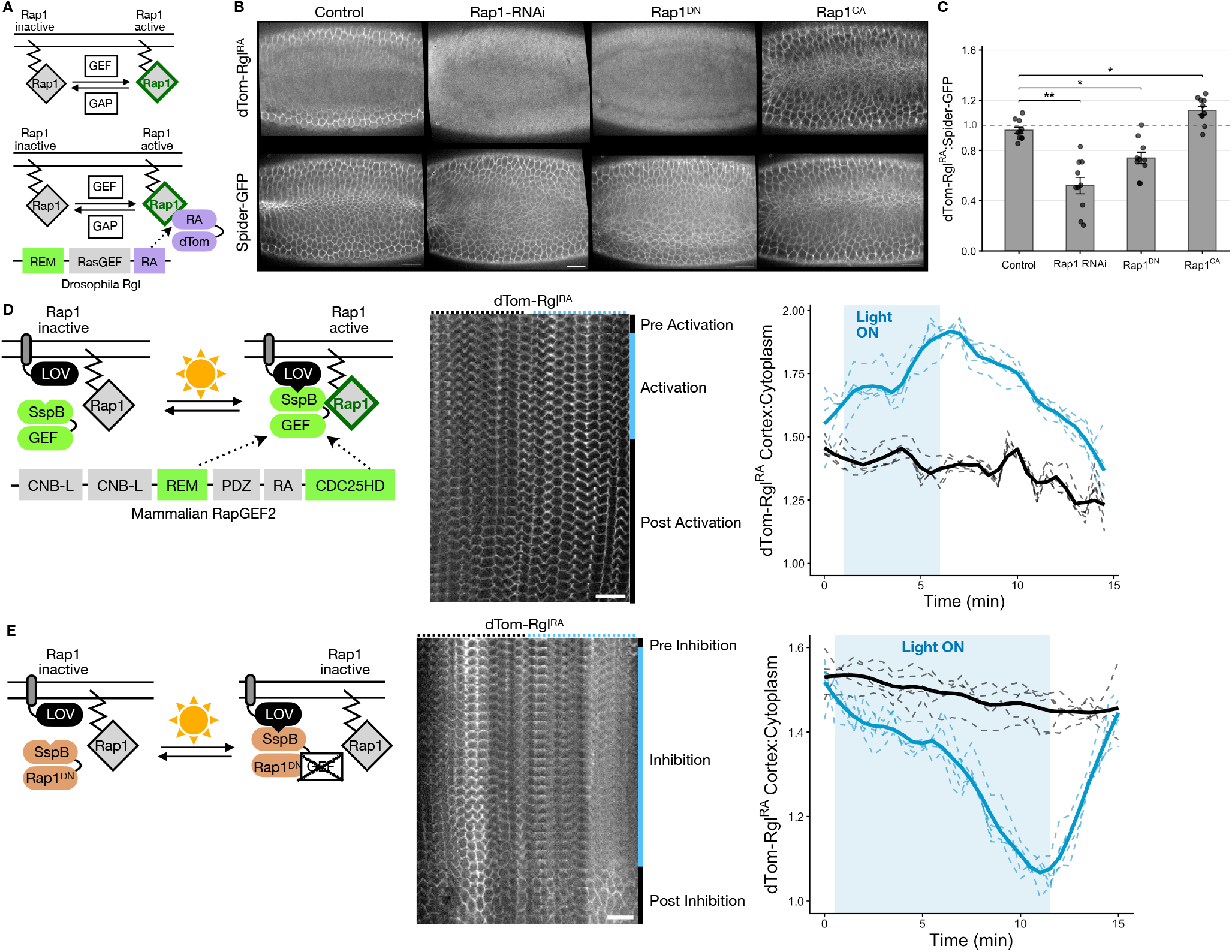
Design and validation of a Rap1 activity biosensor and optogenetic tools for acute manipulation of Rap1 activity *in vivo*. (A) Top, schematic of Rap1 GTPase cycling between inactive GDP-bound and active GTP-bound states. Rap1 activation is promoted by Guanine nucleotide Exchange Factors (GEFs); GTP hydrolysis and consequent Rap1 inactivation are accelerated by GTPase-Activating Proteins (GAPs). Bottom, design of the Rap1 activity biosensor. The Ras-association (RA) domain of *Drosophila* Rgl, which binds active Rap1 preferentially, is fused to dTomato (dTom-Rgl^RA^). Active, membrane-associated Rap1 therefore recruits the biosensor to the cortex, and cortical enrichment reports relative Rap1 activity. Below, domain organization of full-length Rgl: a Ras-exchanger motif (REM), a Ras guanine nucleotide exchange factor domain (RasGEF), and the RA domain. Only the RA domain is used in the biosensor, so the probe lacks exchange-factor activity. (B) Validation of the biosensor using maternal Rap1 perturbations driven by *mat-α*-Gal4. Ventral views of embryos expressing the biosensor (top) and the membrane marker Spider-GFP (bottom) in control embryos and in response to Rap1-RNAi, Rap1^DN^, or Rap1^CA^. Scale bar, 20 µm. (C) Cortex:cytoplasm ratio of dTom-Rgl^RA^ normalized to that of Spider-GFP. Bars, mean; error bars, SEM; points, individual embryos. n = 10 embryos per condition. One-way ANOVA with Dunnett’s test versus control: Rap1-RNAi p < 0.01, Rap1^DN^ p < 0.05, Rap1^CA^ p < 0.05. (D) Optogenetic activation of Rap1. Left, design: a minimal catalytic fragment of mammalian RapGEF2 (REM– CDC25HD) fused to SspB (optoRap1GEF) is recruited by blue light to membrane-tethered LOV–SsrA, activating Rap1 locally. Domain organization of full-length RapGEF2 is shown, with the recruited, concatenated domains indicated in green. Center, montage of a time course of the Rap1 biosensor before, during, and after illumination in a small portion of the embryo spanning illuminated and control regions; illuminated region is indicated at top with a dashed blue line, adjacent non-illuminated region in dashed black line; time runs top to bottom. Right, cortex:cytoplasm ratio of dTom-Rgl^RA^ in the entire illuminated (blue) and equally sized non-illuminated (black) regions of the same embryos. Dashed lines, individual embryos; thick lines, mean; blue shading, light on. Cortical recruitment increases in the illuminated region with no significant change in the non-illuminated region. n = 5 embryos. Scale bar, 20 µm. (E) Optogenetic inhibition of Rap1. Left, design: a cytosolic, CAAX-less dominant-negative Rap1^S17A^ (Rap1^DNΔCAAX^) fused to SspB (optoRap1^DN^) is recruited to the membrane by blue light, where it sequesters endogenous Rap1 GEFs. Center and right, as in D. Cortical recruitment decreases in the illuminated region with no significant change in the non-illuminated region. n = 5 embryos. Scale bar, 20 μm.

To acutely perturb Rap1 activity, we adapted the iLID-based optogenetic system previously applied to the small GTPase Rho1 in *Drosophila* (Rich et al., 2020) (Fig. 2D, E). In this two-component system, a membrane-tethered, light-sensitive LOV domain is fused to the SsrA peptide (bait); upon exposure to blue light, it recruits cytosolic SspB fused to the protein of interest (prey) (Guntas et al., 2015). We fused SspB to a minimal catalytic fragment of mammalian RapGEF2 (Gao et al., 2001, Kuiperij et al., 2003), sufficient to activate Rap1, generating optoRap1GEF. Recruiting optoRap1GEF to the membrane with blue light increases cortical Rap1-biosensor signal in the illuminated region, with no significant change in paired, non-illuminated regions of the same embryos (Fig. 2D). Biosensor signal rises during illumination, peaking shortly after light exposure at ∼6 min, and then decays with a half-time of ∼4 min, recovering partially by 15 min. To inhibit Rap1 activity, we fused SspB to a cytosolic variant of Rap1^S17A^ (Rap1^S17AΔCAAX^), generating optoRap1^DN^, which lacks nucleotide binding ability and sequesters Rap1 GEFs in unproductive complexes (Feig and Cooper, 1988, Feig, 1999). Light-induced membrane recruitment of optoRap1^DN^ produces the reciprocal response, again confined to the illuminated region. Cortical biosensor signal decreases, reaching its lowest value at ∼11 min, and recovers rapidly following stimulation, returning to the level of the paired non-illuminated regions within ∼3 min (Fig. 2E). The reversibility of both perturbations indicates that endogenous Rap1 cycles actively during early embryogenesis, and, the biosensor reports these changes in real time. Together, these reagents allow Rap1 activity to be read out and manipulated with high temporal precision.

### Rap1 activity is reduced in ventral cells during ventral furrow formation

We used the Rap1-biosensor to follow Rap1 activity across successive stages of early embryonic development, surveying the late nuclear cycles of the syncytial embryo, cellularization, and VFF. During nuclear cycles 11–13 and cellularization, the Rap1-biosensor is uniformly enriched at the cortex along the dorsoventral axis (Fig. 3A, Videos S3 and S4). As the new lateral membranes are built, the signal becomes enriched apicolaterally. A separate basal pool is also apparent, although the detection of fluorescent protein attenuates with depth, so we did not analyze this pool quantitatively (Fig. 3—Supplement 1).

**Figure 3.**
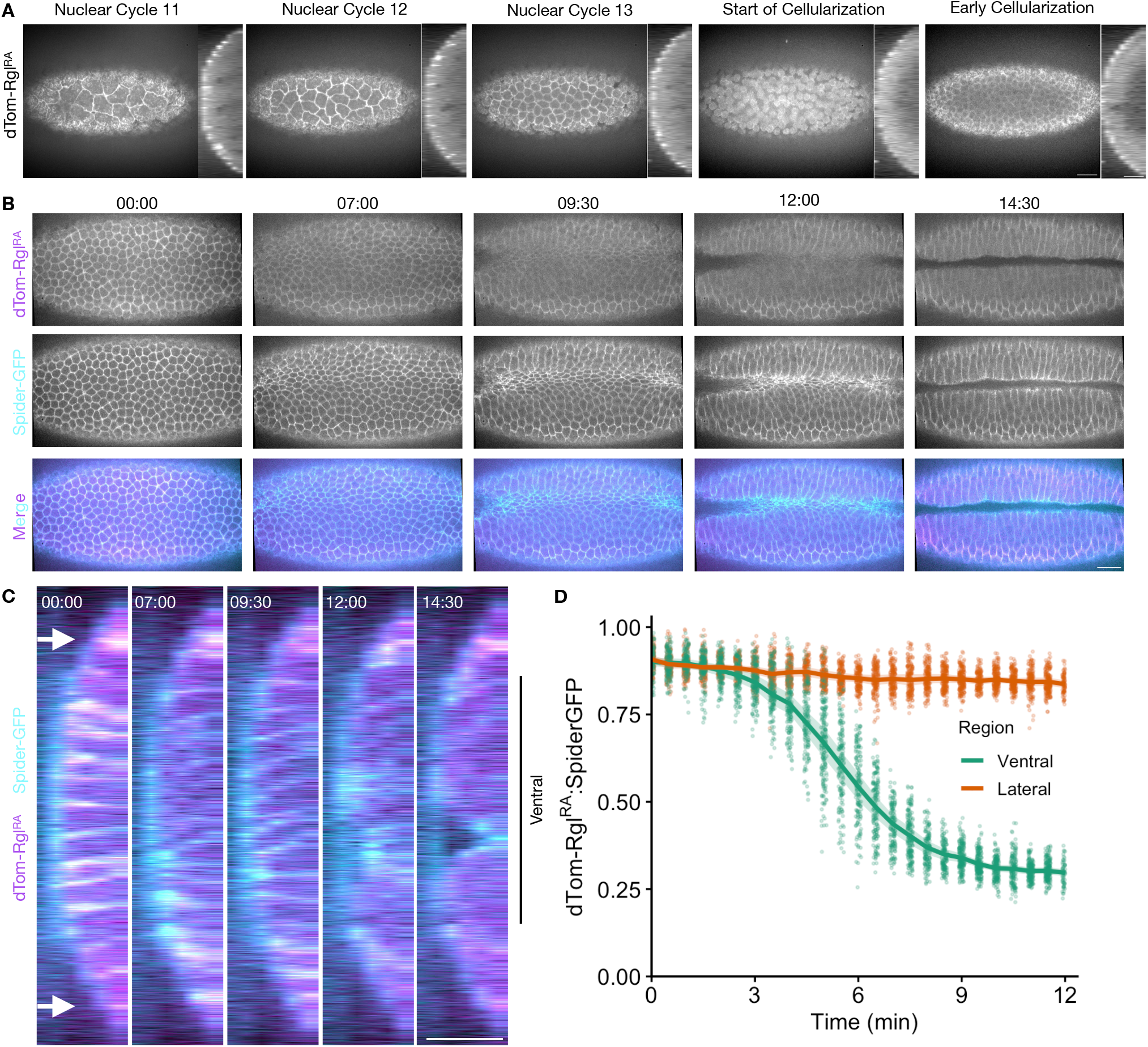
Rap1 activity is reduced in ventral cells during the period of furrow formation. (A) Representative images of embryos expressing the Rap1 biosensor dTom-Rgl^RA^ across nuclear cycles 11– 13, the start of cellularization, and early cellularization. Ventral:lateral cortical biosensor ratio ≈ 1.0 across stages (orthogonal views, right of each frame). n = 5 embryos. Scale bar, 20 μm. (B) *En face* time series during ventral furrow formation in embryos co-expressing dTom-Rgl^RA^ and the membrane marker Spider-GFP, shown as single channels and a merged view. Time (mm:ss), with 00:00 marking the onset of ventral furrow formation. n = 5 embryos. Scale bar, 20 μm. (C) Orthogonal (cross-sectional) views of the same embryos and timepoints as (B), co-expressing dTom-Rgl^RA^ (magenta) and Spider-GFP (cyan). The reduction of cortical dTom-Rgl^RA^ is confined to the ventral domain (bracket on right), while lateral regions (white arrows) retain cortical biosensor enrichment. Scale bar, 20 μm. (Note the straight line at top of embryo in YZ projections reflects autofluorescence from embryo mounting.) (D) Quantification of the cortical biosensor ratio (dTom-Rgl^RA^:Spider-GFP) over time during furrowing, for the ventral (green) and lateral (orange) domains (t = 0, furrow onset; each point, per-cell measurement; line, mean; band, SEM; n = 5 embryos).

The uniformity in Rap1 activity profile is lost during VFF. Beginning a few minutes after the onset of furrowing, cortical biosensor signal decreases in the ventral epithelium, dropping ∼3-fold as furrowing progresses. The signal in the lateral ectodermal cells, in contrast, remains high throughout this period (Fig. 3B–D, Video S5). High Rap1 activity is thus not maintained in ventral cells during furrowing, despite Rap1 being essential for furrow formation.

### Optogenetic inhibition of Rap1 from cellularization onset disrupts VFF

We first validated the efficacy of our optogenetic inhibitory probe optoRap1^DN^ by inhibiting Rap1 during cellularization and VFF, with the ultimate goal of defining the temporal window during which Rap1 is required by inhibiting it during VFF, early cellularization alone, and late cellularization and VFF.

Unlike chronic depletion by RNAi or in germline clones, optogenetic Rap1 inhibition from the onset of cellularization through VFF leaves Rap1 activity in the preceding syncytial nuclear cycles unperturbed (Fig. 4A, Video S6). Inhibition across this time window severely disrupts VFF (Fig. 4B, Video S7). Apical constriction is much slower, with the constriction rate reduced by 74% (Fig. 4C). The constrictions also become progressively less coordinated, with variability in apical area increasing over time while remaining steady in controls (Fig. 4D). Constriction is also markedly less anisotropic, in both magnitude and alignment (Fig. 4E). Thus, optogenetic inhibition of Rap1 beginning at cellularization closely reproduces the phenotype of chronic Rap1 depletion (Asha et al., 1999, Spahn et al., 2012).

**Figure 4.**
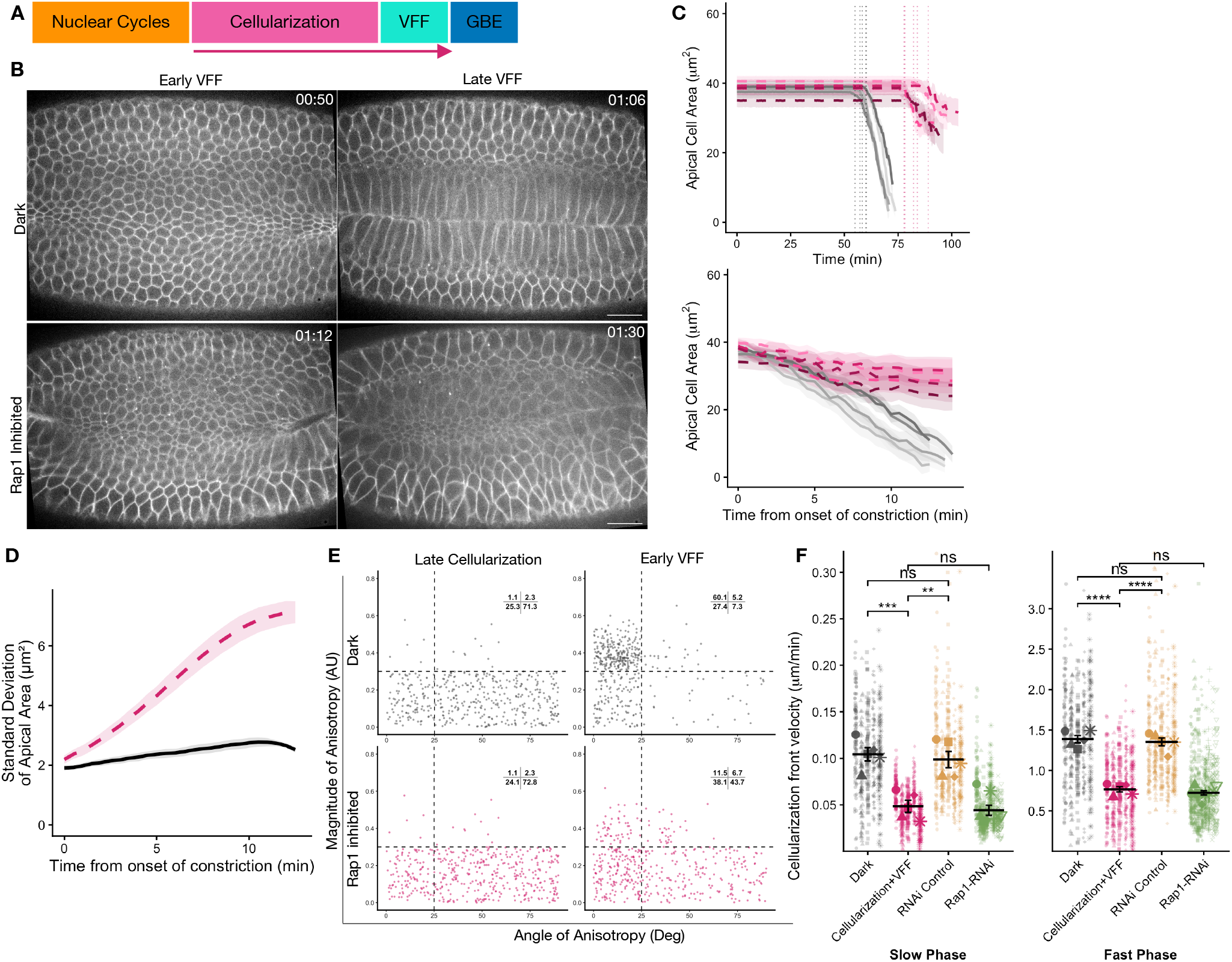
Optogenetic inhibition of Rap1 from cellularization onset through ventral furrowing disrupts ventral furrow formation. (A) Schematic of early embryonic development marking the illumination window (magenta arrow), which begins at the onset of cellularization and continues through ventral furrow formation (VFF). (B) Time-lapse images of optoRap1^DN^ embryos co-expressing Gap43-mCherry, under dark-control conditions (top) or global Rap1 inhibition (bottom). Frames are shown at early and late VFF. Time (hh:mm) relative to the onset of cellularization. Scale bars, 20 µm. (C) Apical area over time in dark-control (gray) and Rap1-inhibited (magenta) embryos. Top, each trace aligned to the onset of cellularization, with dotted vertical lines marking the onset of apical constriction in each embryo, defined as a 5% decrease in apical area. Bottom, the same traces realigned temporally to constriction onset. Each line is the mean apical area of one embryo; shaded bands, SD across cells within that embryo. n = 5 embryos per condition. (D) Standard deviation of apical area across cells within each embryo, over time, in dark-control (black) and Rap1-inhibited (magenta) embryos. Lines, mean; shaded band, SEM; n = 5 embryos per condition. (E) Cell-shape anisotropy at late cellularization (left) and early VFF (right) in dark-control (top) and Rap1-inhibited (bottom) embryos. Each point is one cell, plotted by anisotropy magnitude (arbitrary units) against orientation angle (degrees). Dashed lines partition the plot into quadrants; inset values give the percentage of cells in each. n = 5 embryos per condition. (F) Cellularization front velocity during the slow and fast phases in dark controls (gray), embryos inhibited from cellularization onset through VFF (magenta). RNAi controls (orange), and maternal Rap1-RNAi (green) are replotted for comparison. Horizontal lines and error bars, mean and SEM of embryo means; points, individual front measurements; large symbols, per-embryo means. Statistics are Welch’s t-tests on per-embryo means. N = 5 embryos per condition.

Using this dataset, we further quantified the consequences of this inhibition on the kinetics of cellularization. Inhibition from cellularization onset reduces the rate of front ingression during both the slow and fast phases relative to dark controls (Fig. 4F). These reduced rates are comparable to those of maternal Rap1 RNAi, indicating that this prolonged optogenetic inhibition effectively mimics chronic depletion. RNAi-mediated depletion of maternal Canoe, the Rap1 effector required for apical junction positioning, does not slow cellularization, although these embryos also show ventral furrow defects (Fig. 4—Supplement 1). This suggests that the Rap1 requirement in promoting cellularization is distinct from its role in junction positioning.

Optogenetic inhibition of Rap1 during cellularization also delays the onset of VFF (Fig. 4C). Normally, VFF and germ-band extension overlap only briefly. The two are driven by visibly distinct cell shape changes in different populations of cells: apical constriction of ventral cells drives VFF and oriented cell intercalation in the lateral ectoderm drives the posterior-directed movement during germ-band extension (Bertet et al., 2004, Zallen and Wieschaus, 2004, Blankenship et al., 2006, Stern et al., 2022). We therefore asked whether the delay caused by slow cellularization increases the overlap between these two events. We tracked posterior-directed movement of ventral cells under the same inhibition regime. In dark controls, substantial posterior-directed movement of ventral tissue occurs only after furrow internalization is complete. In Rap1-inhibited embryos, ventral cells are displaced posteriorly while still undergoing the apical constrictions characteristic of early VFF (Fig. 4—Supplement 2). The overlap with germ-band extension is therefore evident during the constriction phase of furrowing. This raises the possibility that orthogonal, posterior-directed forces from timely germ-band extension contribute to the severity of the ventral furrow defect.

Given that actomyosin contractility generates the forces that drive cellularization (Foe and Alberts, 1983), the defects caused by optogenetic Rap1 inhibition might arise from disrupted actomyosin organization. We therefore examined myosin localization during cellularization and VFF. During cellularization, myosin localizes to the tips of the ingressing fronts in both control and Rap1-inhibited embryos, and, in the latter case, this localization persists despite the slower ingression. Later, myosin accumulates apically and forms foci as constrictions become visible, though these events are delayed when Rap1 is inhibited (Fig. 4—Supplement 3). The defects in cellularization therefore do not arise from a failure to accumulate myosin, consistent with reports that cytoskeletal components are correctly localized at the cellularization front upon chronic Rap1 depletion (Spahn et al., 2012, Choi et al., 2013). We cannot, however, rule out a defect in coupling of the actomyosin cytoskeleton to the membrane at the ingressing front. Together, these results demonstrate potent inhibition of Rap1 using these optogenetic tools.

### Acute Rap1 inhibition during furrowing modestly affects VFF

Previously reported chronic Rap1 depletion and our optogenetic inhibition throughout cellularization and VFF both delay and impair VFF. This defective phenotype could result from a direct requirement for Rap1 during furrowing itself (Asha et al., 1999, Spahn et al., 2012) (Fig. 1—Supplement 2). However, this interpretation does not readily align with the reduction in ventral Rap1 activity we observe as the furrow forms (Fig. 3). The phenotype could instead reflect a requirement for Rap1 activity at earlier stages. To test for a Rap1 requirement during furrowing, we acutely inhibited Rap1 only during VFF.

Embryos in which Rap1 is acutely inhibited during VFF form a furrow with morphology similar to that of dark controls (Fig. 5A, B, Video S8). Apical constriction is slightly slowed (constriction rate reduced by 21%; Fig. 5C). Coordination among cells is slightly reduced, with apical cell area becoming more variable as constriction progresses (Fig. 5D). Furrow sealing is less precise, following a more irregular line than the nearly straight seal of controls (Fig. 5— Supplement 1). The anisotropy of apical constriction is unchanged in magnitude and alignment (Fig. 5E). These defects are modest: constriction remains sufficient to drive internalization, and the furrow forms in every case.

**Figure 5.**
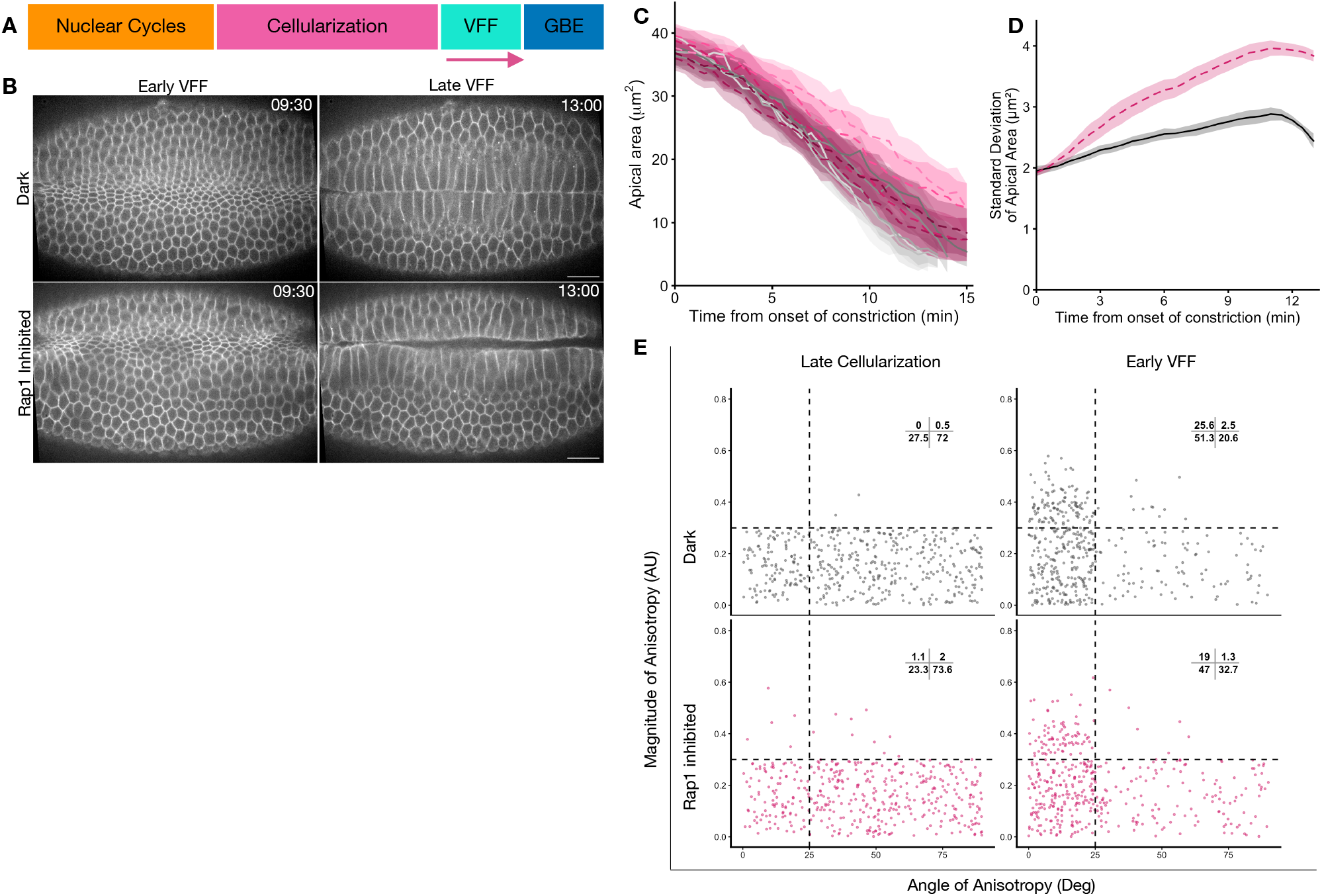
Acute optogenetic inhibition of Rap1 during furrowing does not significantly impair ventral furrow formation. (A) Schematic of early embryonic development marking the illumination window (magenta arrow), which begins at the onset of ventral furrow formation (VFF). (B) Time-lapse images of optoRap1^DN^ embryos co-expressing Gap43-mCherry, under dark-control conditions (top) or global Rap1 inhibition from VFF onset (bottom). Frames are shown at early and late VFF. Time (mm:ss) relative to end of cellularization. Scale bars, 20 µm. (C) Apical cell area over time in dark-control (gray, solid) and Rap1-inhibited (magenta, dashed) embryos, with t = 0 at the end of cellularization. Each line is the mean apical area of one embryo; shaded bands, SD across cells within that embryo. n = 5 embryos per condition. (D) Standard deviation of apical area across cells within each embryo, over time, in dark-control (black) and Rap1-inhibited (magenta) embryos. Lines, mean across embryos; shaded band, SEM; n = 5 embryos per condition. (E) Cell-shape anisotropy in dark-control (top) and Rap1-inhibited (bottom) embryos at late cellularization (left, before inhibition) and early VFF (right, after inhibition and before internalization). Each point is one cell, plotted by anisotropy magnitude (arbitrary units) against orientation angle (degrees). Dashed lines partition the plot into quadrants; inset values give the percentage of cells in each. n = 5 embryos per condition

Rap1 activity is thus not acutely required for the furrow to form and internalize. This is consistent with the reduced ventral Rap1 activity we observe during furrowing (Fig. 3) and contrasts with the strong ventral furrow phenotype of chronic Rap1 depletion (Asha et al., 1999, Spahn et al., 2012) (Fig. 1—Supplement 2) and Rap1 inhibition spanning cellularization and VFF (Fig. 4). The essential requirement for Rap1 therefore lies before the furrowing window, during cellularization. However, Rap1 does appear to play a minor role during VFF. We decided to focus on the more consequential defect resulting from Rap1 inhibition at earlier stages.

### Transient Rap1 inhibition restricted to early cellularization is sufficient to disrupt VFF

Optogenetic Rap1 inhibition during both cellularization and furrowing severely disrupts VFF (Fig. 4), whereas inhibition during furrowing alone produces only modest defects (Fig. 5). These results implicate cellularization as the window during which Rap1 activity is most critical for VFF. To narrow this window, we restricted the perturbation to the early slow phase of cellularization, releasing the inhibition once the cellularization front reached a depth of 10 µm (Fig. 6A). This acute inhibition allows Rap1 activity to recover while the remainder of cellularization completes before furrowing begins. Upon this transient inhibition, the slow-phase rate is reduced by half, comparable to the rate under prolonged inhibition, whereas the fast-phase rate recovers to about 70% of that in dark controls (Fig. 6B; Video S10). The effect of Rap1 inhibition on cellularization kinetics is therefore fast and reversible.

**Figure 6.**
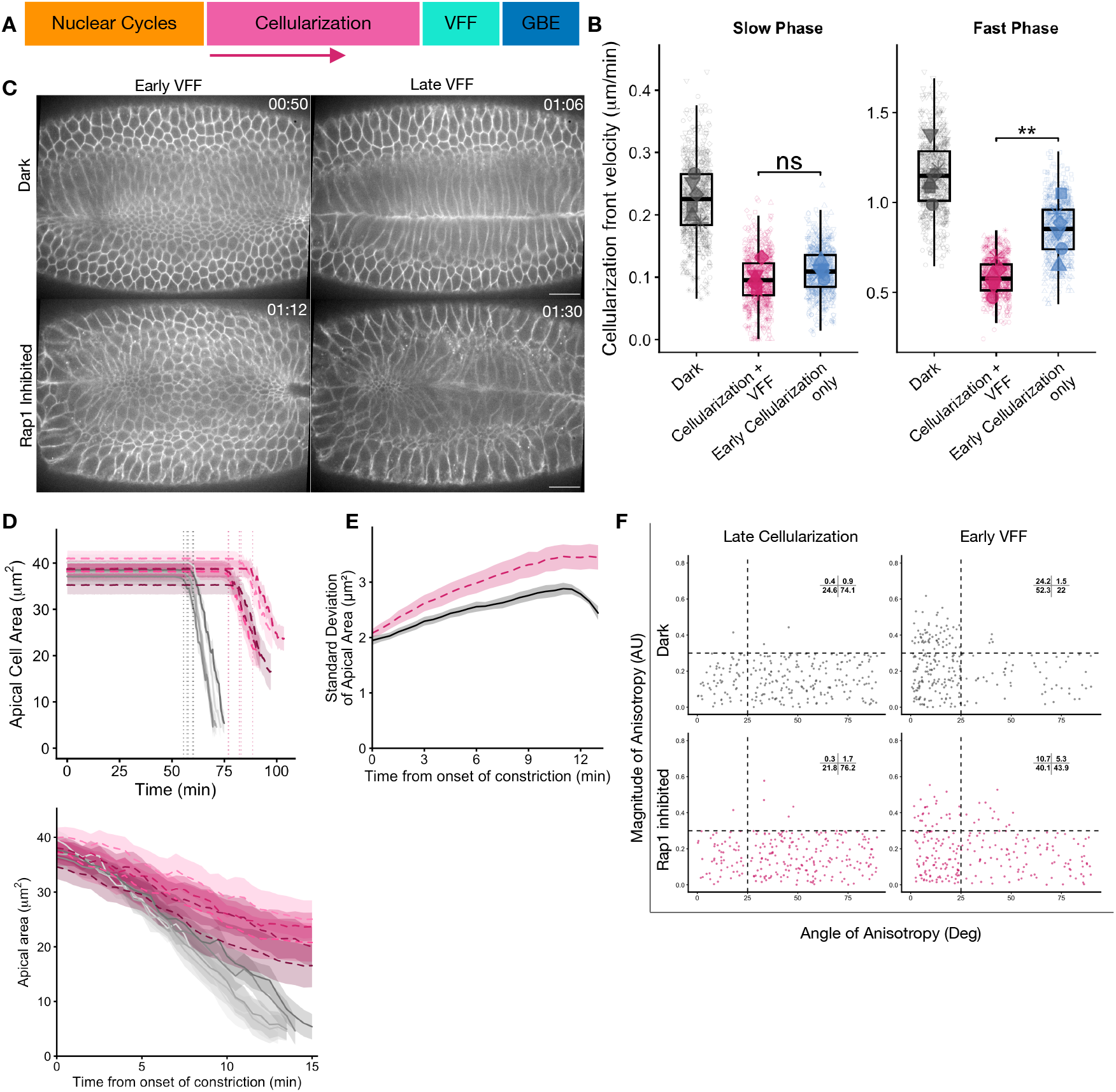
Transient optogenetic Rap1 inhibition during early cellularization is sufficient to disrupt VFF. (A) Schematic of early embryonic development marking the illumination window (magenta arrow), which spans the early slow phase of cellularization and ends when the cellularization front reaches a depth of 10 µm. (B) Cellularization front velocity during the slow and fast phases in dark controls (gray), embryos inhibited throughout cellularization and VFF (magenta), and embryos inhibited during early cellularization only (blue). Boxes, median and IQR; small symbols, individual front measurements, with shape denoting the embryo of origin within each condition; large symbols, per-embryo means. Welch’s t-tests on per-embryo means, n = 5 embryos per condition. (C) Time-lapse images of optoRap1^DN^ embryos co-expressing Gap43-mCherry, under dark-control conditions (top) or Rap1 inhibition restricted to early cellularization (bottom). Frames are shown at early and late VFF. Time (hh:mm) relative to the end of nuclear cycle 13. Scale bars, 20 µm. (D) Apical cell area over time in dark-control (gray, solid) and Rap1-inhibited (magenta, dashed) embryos. Top, traces aligned to the onset of cellularization; dotted vertical lines mark the onset of apical constriction in each embryo, colored by condition and defined as a 5% decrease in apical area. Bottom, the same traces realigned to constriction onset. Each line is the mean apical area of one embryo; shaded bands, SD across cells within that embryo. n = 5 embryos per condition. (E) Standard deviation of apical area across cells within each embryo, over time, in dark-control (black) and Rap1-inhibited (magenta) embryos. Lines, mean across embryos; shaded band, SEM; n = 5 embryos per condition. (F) Cell-shape anisotropy in dark-control (top) and Rap1-inhibited (bottom) embryos at late cellularization (left) and early VFF (right). Each point is one cell, plotted by anisotropy magnitude (arbitrary units) against orientation angle (degrees). Dashed lines partition the plot into quadrants; inset values give the percentage of cells in each. n = 5 embryos per condition.

Despite the recovery of Rap1 activity before furrowing, VFF is highly aberrant. Acute inhibition during early cellularization slows apical constriction, as does the prolonged inhibition described above (Fig. 6C). Apical area becomes increasingly variable as constriction proceeds, suggesting decreased coordination among cells (Fig. 6D). Transient inhibition also reduces constriction anisotropy in both magnitude and alignment (Fig. 6E). Slowed cellularization resulting from this transient inhibition delays furrow onset, so that germ-band extension begins while ventral cells are still constricting (Fig. 6—Supplement 1). Together, these observations indicate that active Rap1 is required during the early, slow phase of cellularization for normal VFF, since restoring activity before furrowing does not rescue these defects.

When Rap1 inhibition is restricted to the early slow phase, myosin remains associated with the ingressing front despite the slower ingression and subsequently accumulates apically into foci coincident with the appearance of constrictions, as in dark controls (Fig. 6—Supplement 2; Video S11-13). This matches the behavior seen when optogenetic inhibition spans both cellularization and furrowing. The defects therefore do not arise from a failure to localize or accumulate myosin in ventral cells.

To test whether this requirement is specific to early cellularization, we inhibited Rap1 acutely during late cellularization and VFF (Fig. 6—Supplement 3). These embryos form a ventral furrow with only modest defects, comparable to those seen upon inhibition during furrowing alone (Fig. 4). Because this perturbation lasts longer than inhibition during furrowing alone, it also rules out insufficient Rap1 inhibition as an explanation for the mild phenotype. The fast-phase ingression rate is slowed, but due to its short duration, the resulting delay does not produce the extensive overlap with germ-band extension seen upon inhibition during early cellularization. Rap1 activity is therefore largely dispensable during late cellularization and VFF.

Together, the four perturbations narrow the temporal requirement of Rap1 in VFF: inhibition confined to the early slow phase of cellularization is sufficient to disrupt furrowing, whereas inhibition during late cellularization and/or furrowing leaves the process largely intact (Fig. 5; Fig. 6—Supplement 3). Rap1 activity therefore acts within a discrete critical window to ensure successful VFF. The requirement for cellularization kinetics, however, is markedly different: ingression slows when Rap1 is inhibited and recovers when activity is restored, indicating a continuous requirement throughout ingression rather than in a restricted window.

### Early Rap1 activity is required for adherens junction establishment

Adherens junctions assemble progressively during cellularization (Harris and Tepass, 2010). At the outset of cellularization, when the embryo is still a syncytium, cadherin-catenin complexes are present at the membrane but have not yet clustered into adherens junctions (Hunter and Wieschaus, 2000, Harris and Peifer, 2004). As the lateral membrane elongates, these complexes enrich at the apicolateral position and coalesce into spot adherens junctions. The spots later mature into a continuous belt at the beginning of gastrulation (Müller and Wieschaus, 1996, Hunter and Wieschaus, 2000, Harris and Peifer, 2004, Harris and Peifer, 2005). The mature junctions are then maintained and remodeled during VFF, as cells come under tension (Sawyer et al., 2009, Martin et al., 2010). The critical window for Rap1 activity that we identified coincides with the initiation of this assembly during the early slow phase of cellularization. Rap1 may therefore act during this window to enable assembly of the apical junctions that are required later during furrowing, consistent with previous suggestions (Choi et al., 2013).

By combining optoRap1^DN^ with an E-cad:GFP reporter, we asked whether inhibiting Rap1 activity during cellularization affects assembly of the adherens junctions. Although optoRap1^DN^ is also GFP-tagged, its cortical intensity is negligible relative to that of E-cad:GFP, so the cortical signal we measure reports E-cadherin (optogenetics-compatible E-cad reporters such as E-cad:mKate2, E-cad:tagRFP and E-cad:tdTomato are not detectable at these stages). Because blue light both excites E-cad:GFP and recruits optoRap1^DN^ to the membrane, inhibition cannot be separated from imaging, precluding an inhibition-and-recovery protocol. We therefore used continuous inhibition (as in Fig. 5A). In control embryos expressing E-cad:GFP only (without optoRap1^DN^), we observe two pools of E-cadherin (Fig. 7A): an apicolateral pool that persists into VFF, and a transient basal pool that tracks the ingressing front and is lost at the onset of furrowing. Both match the previously described patterns (Hunter and Wieschaus, 2000).

**Figure 7.**
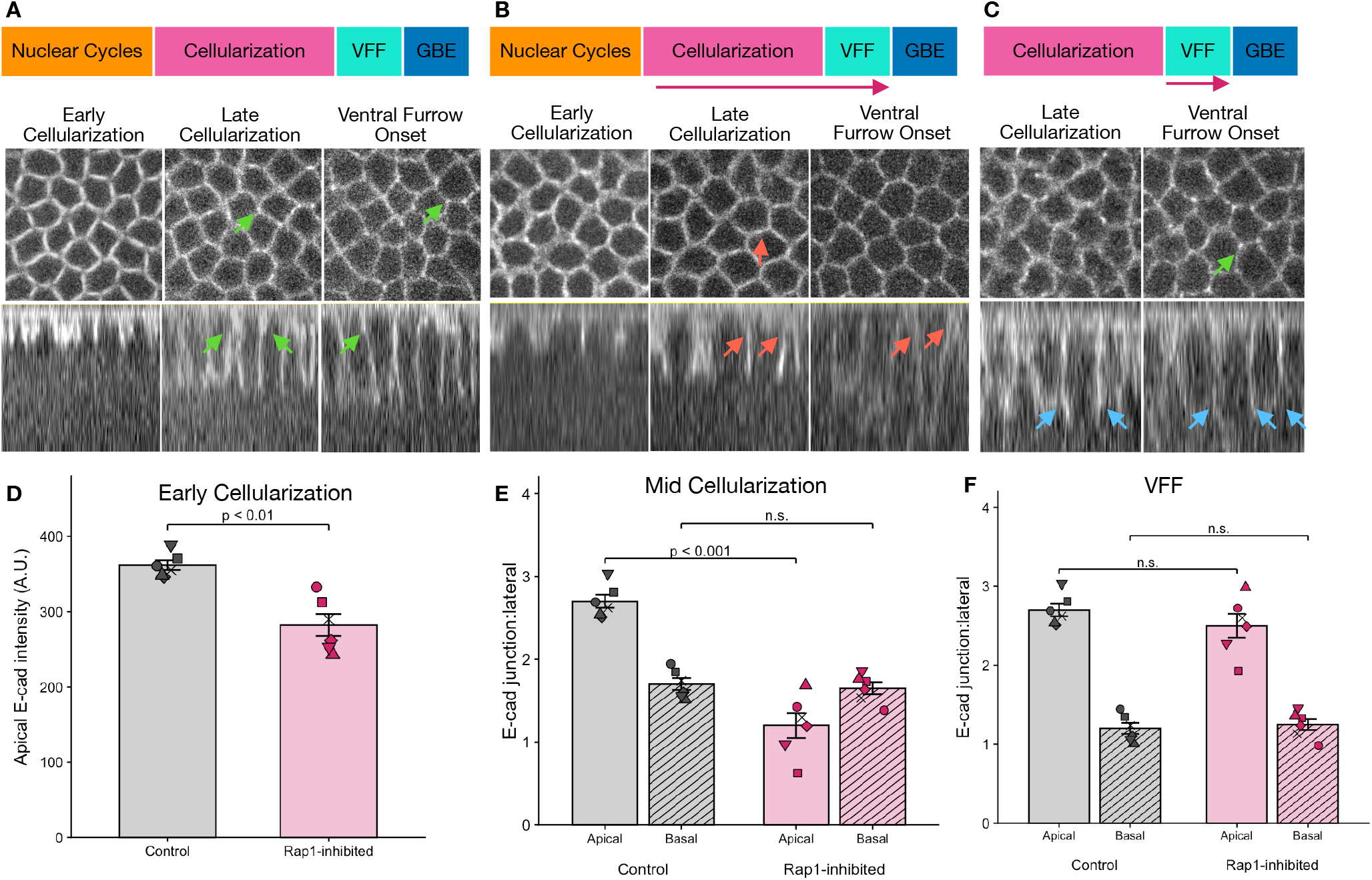
Rap1 activity during early cellularization is required for adherens junction establishment. (A) E-cadherin localization in control embryos at early cellularization, late cellularization, and ventral furrow onset, shown en face (top) and in orthogonal (XZ) views (bottom). E-cadherin becomes progressively enriched at the apicolateral membrane and at tricellular junctions as cellularization proceeds (green arrows). Schematic (top) indicates the developmental window imaged. (B) E-cadherin localization when Rap1 is optogenetically inhibited from the onset of cellularization (inhibition window indicated by magenta arrow in the top schematic). Apical E-cadherin enrichment fails to develop, and predominantly basal signal remains by late cellularization (red arrows); the loss persists to ventral furrow onset. (C) E-cadherin localization when Rap1 is inhibited only during ventral furrow formation, after cellularization is complete (inhibition window indicated by magenta arrow in the top schematic). Apical and tricellular E-cadherin enrichment established during cellularization is preserved (green arrows). Blue arrows, basal junctions, which disappear at ventral furrow onset. (D) Background-subtracted E-cadherin intensity in control (gray) and Rap1-inhibited (magenta) embryos at early cellularization. (E) Junctional E-cadherin intensity normalized to lateral E-cadherin intensity in control (gray) and Rap1-inhibited (magenta) embryos at mid cellularization when fronts have reached a depth of 20 μm. Apical junctions in solid bars and basal junctions in dashed bars. (F) Junctional E-cadherin intensity normalized to lateral E-cadherin intensity in control (gray) and Rap1-inhibited (magenta) embryos during VFF. Apical junctions in solid bars and basal junctions in dashed bars. Arrows: green, sites of normal apical/tricellular E-cadherin enrichment; red, sites of failed enrichment. n = 5 embryos per condition. Welch’s t-tests on per-embryo means (D, E, F).

**Figure 8.**
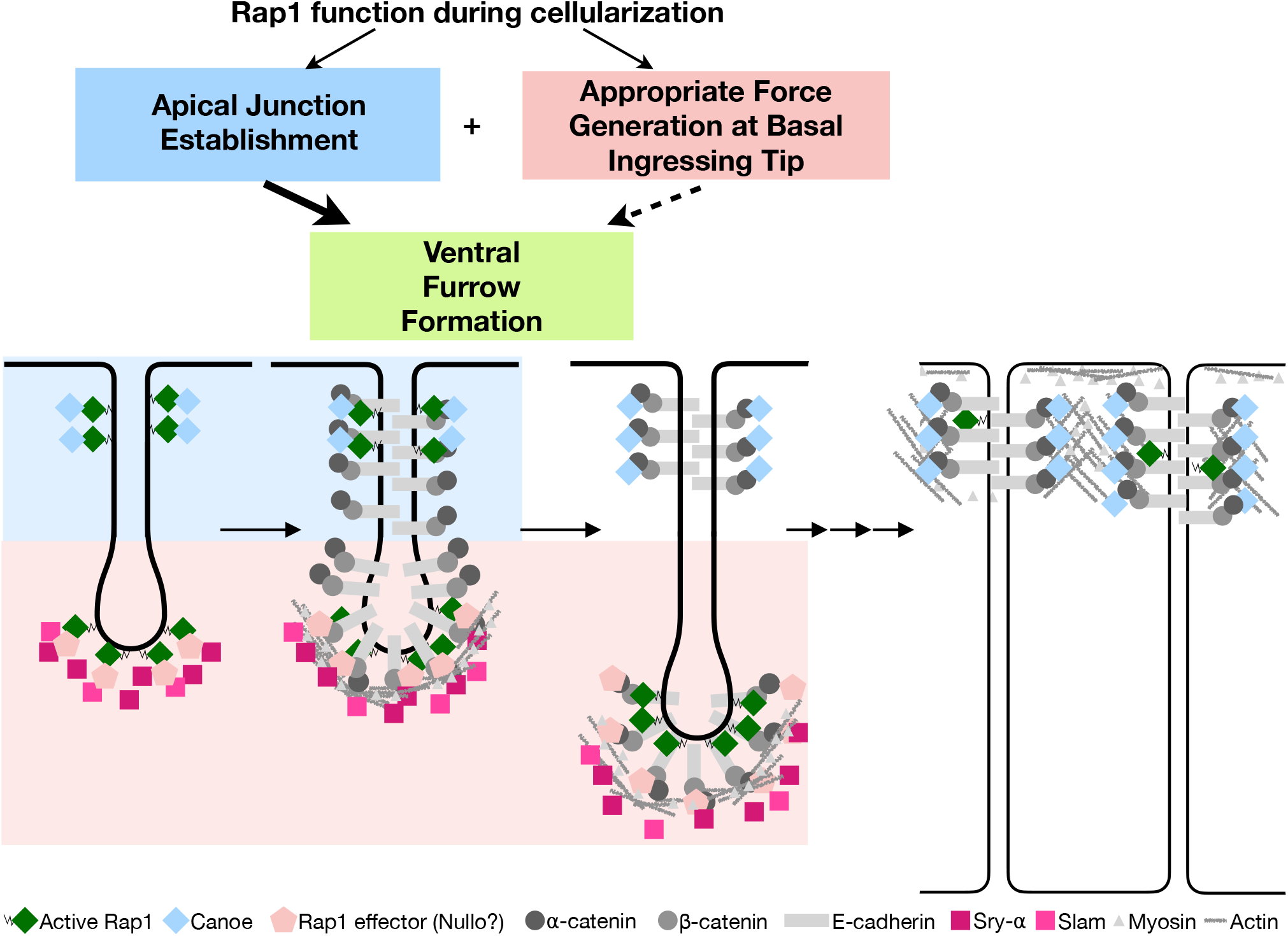
Model for Rap1 function during cellularization and its consequences for ventral furrow formation. Top. The two separable Rap1 functions identified here and their contributions to ventral furrow formation (VFF). During the early slow phase of cellularization, active Rap1 acting through its effector Canoe establishes apical adherens junctions, a step that also requires Bazooka and that depends on the Rap1 GEF ELMO-Sponge (Left, blue). Throughout cellularization, Rap1 is required for membrane ingression to proceed at an appropriate rate. We propose that this reflects linkage of basal junctions to the actomyosin cytoskeleton through an effector that remains unidentified (possibly Nullo), and that slowed ingression retains myosin at the basal tip, delaying its availability at the medioapical cortex and thereby the onset of apical constriction (Right, pink). Successful VFF critically requires the apical junctions established during early cellularization. At the onset of furrowing, Rap1 activity falls in ventral cells and myosin is released from the basal tip, coincident with Nullo degradation (green). Solid arrows, relationships supported by the data presented here; weighted arrow, primary VFF requirement; dashed arrow, proposed. Bottom. Schematic of the molecular events at a single cellularization front, apical up, at four successive stages: early cellularization, early-to-mid cellularization, late cellularization, and VFF. Blue shading, the apicolateral region, where Rap1 activity is required transiently to establish adherens junctions; pink shading, the basal ingressing tip, where Rap1 activity is required continuously. Active Rap1 is present in both compartments but acts through different effectors: Canoe apicolaterally, and an unidentified effector basally (likely Nullo). See key for symbols.

We quantified E-cadherin at early and late stages of cellularization. At the earliest stages, before a lateral membrane has formed, we compared background-subtracted E-cadherin intensity directly between conditions, as no other membrane region is available for normalization (Fig. 7D). At later stages, a lateral membrane is available for normalization, but its length differs between conditions because Rap1 inhibition slows ingression. We therefore compared control and Rap1-inhibited embryos at matched ingression depths. We quantified apical (or basal) enrichment as the ratio of E-cadherin intensity in an apical (or basal) window to that at a more lateral position (Fig. 7E, F).

Under continuous Rap1 inhibition, less E-cadherin accumulates at the nascent apicolateral membrane at the earliest stages of cellularization (Fig. 7A, B left panel and quantification in 7D). By mid-cellularization (20 µm ingression depth), when controls show clear apicolateral enrichment, apical E-cadherin remains lower in Rap1-inhibited embryos. The basal pool, in contrast, is unaffected and reaches intensities comparable to controls. Rap1 inhibition therefore impairs apical junction assembly without affecting accumulation of the basal pool of E-cadherin (Fig. 7A, B center panel and quantification in 7E). At VFF onset, the apical junctions are enriched for E-cadherin and the basal junctions disappear in control embryos. In contrast, in Rap1-inhibited embryos, while the basal junctions disappear as in controls, the apical junction pool is not enriched for E-cadherin (Fig. 7A, B right panel). Inhibiting Rap1 only during VFF, by contrast, does not disrupt the apical E-cadherin enrichment established during cellularization (Fig. 7C and quantification in 7F). The same stage dependence holds at tricellular junctions, which are sites of elevated tension during morphogenesis (Letizia et al., 2019, Yu and Zallen, 2020). E-cadherin enrichment at tricellular vertices is lost when Rap1 is inhibited during cellularization but retained when inhibition is restricted to furrowing (Fig. 7A-C). Rap1 activity is therefore required to establish apical and tricellular junctions during early cellularization but not for their maintenance during furrowing.

## Discussion

Rap1 is a conserved small GTPase that regulates cell adhesion, among other cellular processes, and is essential for morphogenesis (Knox and Brown, 2002, Boettner et al., 2003, Jaśkiewicz et al., 2018). Rap1 contributes to many processes, which complicates the interpretation of its mutant phenotypes. Here we present tools to visualize Rap1 activity directly and to perturb it acutely and reversibly. Using these tools, we identify a critical window of Rap1 activity during cellularization that is required for successful ventral furrow formation. Rap1 activity is largely dispensable during furrowing itself, consistent with our observation that endogenous Rap1 activity declines in ventral cells as the furrow forms. We also find that Rap1 activity affects the rate of cellularization, a function that impacts the temporal separation of ventral furrowing and germ-band extension. This contribution of Rap1 activity in cellularization may have been missed in prior studies which largely relied on analyses of fixed embryos.

### A critical role of Rap1 during early cellularization in apical adherens junction establishment

Rap1 has long been known to be essential for ventral furrowing (Asha et al., 1999, Spahn et al., 2012). Our results narrow the primary requirement for Rap1 to the early, slow phase of cellularization. During this window, active Rap1 drives enrichment of adherens junction components at the apicolateral membrane through its effector Canoe (Choi et al., 2013, Schmidt et al., 2018). These early-established junctions, and the tricellular junctions in particular, are positioned to sustain mechanical coupling and epithelial integrity as actomyosin contractility drives the shape changes that internalize the ventral epithelium.

Inhibiting Rap1 during cellularization reduces apical E-cadherin enrichment and prevents its accumulation at tricellular vertices. By contrast, inhibition restricted to late cellularization and furrowing leaves the prior enrichment intact. Rap1 activity is therefore required to establish apical junctions but not to maintain them once formed, and late Rap1 activity cannot substitute for activity during the early window. Combining these results with prior work, we propose that Rap1 triggers junction assembly at the apicolateral position through Canoe (Choi et al., 2013), after which junctions are maintained without further Rap1 input. Consistent with this model, a Canoe mutant lacking the Rap1-binding RA domains can accumulate at apical junctions when wild-type Canoe is present, indicating that Canoe can be recruited to adherens junctions without binding Rap1 directly (Bonello et al., 2018). Visualizing Canoe during acute Rap1 inhibition would distinguish whether Rap1 activity is required only to recruit Canoe or also to retain it, but this is not currently feasible in live embryos due to a lack of a tagged Canoe compatible with our optogenetic system. Junctions that fail to form during early cellularization do not appear to reach the functional state required for furrowing. Consistent with this interpretation, although apical spot junction proteins accumulate before furrow onset in *rap1* germline clones through Rap1-independent mechanisms, these embryos still fail to furrow (Choi et al., 2013).

This early critical window for junction nucleation may be set by the coincident establishment of apicobasal polarity in the nascent lateral membrane. Bazooka, which marks the apical domain, is required for assembly of apical cadherin complexes at this stage (Harris and Peifer, 2004) and is itself positioned by Rap1 and Canoe (Choi et al., 2013). At the same time, Scribble and Discs-large mark the lateral domain and are also required for the initial positioning and assembly of junctions (Bonello et al., 2019). This partitioning occurs once as the lateral membrane forms. While the cellularization-specific protein Slam is not a polarity marker, it also localizes to the basal front and recruits RhoGEF2 for generating actomyosin contractility at the ingressing front (Wenzl et al., 2010, Schmidt et al., 2018). The junction assembly step that Rap1 acts on may therefore be available only while this polarity partitioning is underway.

### Rap1 activity is also necessary for correctly pacing cellularization

To our knowledge, this is the first report that Rap1 activity is required to sustain the normal rate of cellularization. Our experiments further indicate that the roles of Rap1 in membrane ingression and junction positioning are separable. Rap1 inhibition during late cellularization affects only the rate of membrane ingression and not ventral furrowing, while inhibition during early cellularization affects both. Further, depleting Canoe, the Rap1 effector necessary for junction positioning, does not affect the rate of cellularization but does disrupt furrowing. This experiment also shows that ventral furrow formation depends more on junction establishment than on normal cellularization kinetics. Indeed, embryos that fail to cellularize can nevertheless undergo furrow invagination (Goldner et al., 2024). However, timely cellularization determines whether furrowing begins on schedule.

The forces that drive cellularization are generated, at least in part, at the basal tip of the ingression front (Wieschaus and Sweeton, 1988, Sokac et al., 2023). Slam, Nullo, and Serendipity-α, among the zygotically expressed cellularization-specific proteins, localize to this tip (Postner and Wieschaus, 1994, Lecuit et al., 2002, Zheng et al., 2013). Slam recruits RhoGEF2 to activate Rho1 that drives actomyosin accumulation and promotes membrane ingression (Wenzl et al., 2010). In addition to these proteins, a basal pool of junctional proteins tracks the ingressing tip. Our experiments demonstrate that neither Rap1 activity nor Canoe accumulation is required for the localization or abundance of this basal pool of E-cadherin.

These observations do not, however, rule out a requirement for Rap1 in linking actomyosin to the membrane, plausibly through basal junction proteins. Rap1 recruits cytoskeleton-junction linker proteins in various contexts, and could perform a similar function at this site (Boettner et al., 2003, Sawyer et al., 2009, Yang et al., 2014, Zhang et al., 2014, Plak et al., 2016, Bromberger et al., 2018). The two Rap1 requirements during cellularization differ in kind. Apical junctions require Rap1 activity once, during establishment, and are refractory to its loss thereafter. Ingression, by contrast, slows within minutes of Rap1 inhibition and recovers when its activity recovers, indicating that Rap1 activity is required continuously at the front.

At the cellularization front, Rap1 may function in coordination with the aforementioned cellularization-specific proteins. Specifically, the zygotic protein Nullo is required for basal junction formation. Intriguingly, Nullo overexpression antagonizes assembly of the apical adherens junctions and results in a failure in VFF (Hunter and Wieschaus, 2000, Hunter et al., 2002). These phenotypes resemble those resulting from Rap1 inhibition. We speculate that active Rap1 interacts with Nullo at basal junctions and its overexpression could outcompete Canoe for Rap1 at apical junctions, thereby inhibiting VFF.

Slowed cellularization due to Rap1 inhibition may also affect the availability of myosin for subsequent processes. Myosin appears to be present in limited quantity, and is retained at the basal cellularization fronts when Rap1 is inhibited. This delayed release of myosin could delay its arrival at the medioapical cortex, where it generates the force for the apical constriction that drives furrow formation. Consistent with this, ectopic Rho1 activation during cellularization using optogenetics produces mild apical constrictions as compared to those observed moments later when cellularization is complete (Rich, Nayak, & Glotzer, unpublished observations). Disassembly of basal junctions may promote myosin release. Nullo is degraded during late cellularization; at gastrulation onset the basal junctions are disassembled as myosin relocalizes from the basal tip to the apical cortex (Hunter and Wieschaus, 2000). If Rap1 inhibition extends the time window of myosin association with a slowly ingressing front, its redistribution to the apical cortex would be correspondingly delayed.

The delayed onset of constrictions, and consequently furrowing, causes an overlap of ventral furrowing with germ-band extension, which generates posterior-directed forces that are orthogonal to those driving furrowing. This temporal overlap between the two processes might exacerbate the ventral furrow phenotype, and in milder cases produce twisting of the furrowed epithelium (Spahn et al., 2012). Our results indicate that the delay originates not only from ineffective constrictions as seen in *dizzy* germline clones, but also due to slow cellularization, which may further limit the actomyosin available for constriction.

While Rap1 functions with Canoe in VFF, promoting cellularization front ingression is among the few Canoe-independent Rap1 functions during cellularization. Another such example is the maintenance of uniform cross-sectional cell areas along the apicobasal axis, which is disrupted in *rap1* but not *cno* mutants (Choi et al., 2013). This phenotype might well be another manifestation of delayed cellularization.

## Conclusion

Ventral furrow formation depends on Rap1 activity at specific times and subcellular sites. During the early slow phase of cellularization, Rap1 positions adherens junction components apicolaterally, where they cluster and mature while the nascent membrane is partitioned into apicobasal domains. In parallel, continuous Rap1 activity drives membrane ingression at the basal front, so that the epithelium is cellularized on schedule and the furrow forms within its normal window, before germ-band extension is underway. Neither requirement extends into furrowing itself, when ventral Rap1 activity declines and acute inhibition of Rap1 causes only mild defects. Thus, VFF is largely independent of Rap1 activity during folding but acutely dependent on its activity during the preceding stage of apical junction establishment. The requirement for Rap1 in furrowing is therefore inherited from cellularization, an illustration of how the success of one morphogenetic stage is set by the state the previous stage leaves behind.

## Materials and Methods

### Plasmids

Plasmids used in this study are listed in Supplementary File 1. pUbi-stop-mCD8GFP, containing an attB site, was a gift from T. Lecuit. pUbi>Stargazin-GFP*-LOVSsrA (* denotes silenced GFP) was described previously (Rich et al., 2020). RalGDS-RBD and HA-RapGEF2 were obtained from Addgene (118315, 110162). Plasmids generated in this study were constructed by one-step isothermal *in vitro* recombination (Gibson et al., 2009) or by ligation. pUbi-stop-mCD8GFP was linearized by restriction digestion to excise the stop-mCD8GFP cassette while retaining the Ubi promoter, attB site, 3′UTR and SV40 terminator. Inserts were amplified with primers carrying ≥20-bp overlap to the linearized vector — from plasmid templates, from genomic DNA of *w^1118^* adults (Rgl^RA^, which is intronless) or from genomic DNA of the UAS>Rap1^S17A^ line (Ellis et al., 2013) (optoRap1^DN^) — and assembled into the digested backbone.

### Fly stocks

*Drosophila melanogaster* was cultured using standard techniques at 25°C. Both male and female animals were used. Stocks used in this study include: *pSqh>Gap43-mCherry/TM3*, generated by P-element insertion and was a gift from A. Martin; *pSqh>Sqh-mCherry* (Martin et al., 2009); UAS>Rap1^S17A^ and UAS>Rap1^Q63E^ (Ellis et al., 2013); *gish*>Spider-GFP (BDSC 59025); Jupiter-GFP (BDSC 6836); Ubi>E-cad:GFP (Kyoto Stock Center #109007; (Oda and Tsukita, 2001)); P(mat-tub-Gal4)mat67 (BDSC 7062); UAS>Rap1-RNAi (B:SC 57851); UAS>Canoe-RNAi (VDRC GD 7769; (Dietzl et al., 2007)); Ubi>Stargazin-GFP-LOV-SsrA (attP2) (generated in (Rich et al., 2020)). Control embryos for RNAi experiments carried the driver and markers without an RNAi transgene (P(mat-tub-GAL4)mat67; Gap43-mCherry).

Transgenic flies were generated by PhiC31-directed integration (GenetiVision). Transgenic lines generated for this study include: Ubi>dTom-Rgl^RA^ (VK31); Ubi>GFP-RalGDS^RA^ (VK31); Ubi>SspB-GFP-Rap1^S17AΔCAAX^ (optoRap1^DN^; VK37); Ubi>SspB-RapGEF2(REM-CDC25HD) (optoRap1GEF; VK37). Genotypes used in each experiment are listed in Supplementary File 2.

### Preparation of *Drosophila* embryos for live imaging

Embryos were collected at 25 °C on apple juice agar plates supplemented with yeast paste. Embryos were collected for 60 min and then aged for 30–90 min at 25°C, such that the majority were either entering cortical nuclear cycles (NC 10-13) or late cellularization at the time of mounting. Embryos were dechorionated in 30% bleach for 1 min, rinsed thoroughly in water, aligned on an apple juice agar pad and mounted ventral side down on a coverslip coated with embryo glue (adhesive from double-sided tape dissolved in heptane, allowed to evaporate before use). The coverslip was affixed with petroleum jelly over the central hole of a metal slide. Embryos were covered with halocarbon oil 200 immediately after mounting and were not compressed. All handling of optogenetic lines was carried out under red safelight illumination to avoid premature perturbation.

### Live imaging and optogenetic experiments

Time-lapse recordings of embryos were acquired on a Zeiss Axiovert 200M equipped with a Yokogawa CSU-X1 spinning-disk unit (Solamere), 488 and 561 nm lasers (50 mW; Coherent) and a Prime 95B camera (Photometrics), using a 40×/1.0 numerical aperture (NA) oil objective and µManager software (Edelstein et al., 2014). Global optogenetic perturbation was accomplished by illuminating the entire field with white LED light at 500 ms exposure through the entire Z stack using a Digital Micromirror Device (DMD) projector. Dark controls were embryos of the same genotype imaged with 561 nm illumination only. Because 488 nm light both excites GFP-tagged reporters and recruits SspB-tagged optoRap1GEF or optoRap1^DN^ to the membrane, experiments using GFP reporters were performed under continuous illumination and could not be combined with an inhibition-and-recovery protocol; for these experiments, controls were genotype-matched embryos lacking the SspB transgene, imaged with identical settings. Illumination protocols for each experiment are listed in Supplementary File 3.

### Image analysis and Statistics

All images were processed and analyzed in Fiji/ImageJ v1.54p (Schindelin et al., 2012). Statistical analysis and plotting were performed in R v4.5.1 using RStudio. Because measurements of individual cells and of the cellularization front are nested within embryos, the embryo was treated as the unit of replication throughout: per-embryo means were computed, and comparisons between conditions were made with Welch’s t-tests unless otherwise noted. Quantification pertaining to each experiment including sample sizes, test statistics and p values are detailed in the Supplementary file 4.

### Structural comparison of RA domains

The RA domain of human RALGDS was taken from the solution NMR ensemble PDB 1RAX. No experimental structure of *Drosophila melanogaster* Rgl (UniProt Q9U4U5) is available, so the AlphaFold DB model AF-Q9U4U5-F1 (Jumper et al., 2021, Varadi et al., 2024) was used and the RA domain, residues 849–948 by UniProt annotation, extracted. Residue correspondence between the two domains was established with TM-align (Zhang and Skolnick, 2005).

## Acknowledgements

This work was supported by R35 GM127091 from NIGMS at the National Institutes of Health. We thank Richard Fehon and Ashley Rich for helpful comments on this manuscript. We thank Ed Munro, Richard Fehon, Elizabeth Heckscher, Glotzer lab members, and Fehon lab members for helpful discussions and support. Stocks obtained from the Bloomington Drosophila Stock Center (NIH P40OD018537) were used in this study.

## Figure and Video Legends

**Figure 1—Supplement 1.**
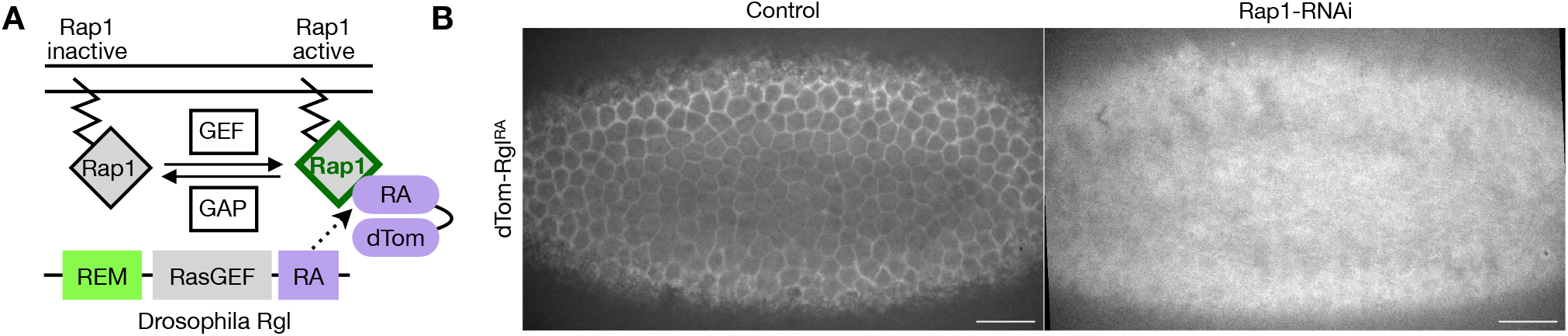
Maternal Rap1-RNAi reduces cortical Rap1 biosensor signal. (A) Schematic of the Rap1 activity biosensor. The Ras-Association (RA) domain of Drosophila Rgl, which binds GTP-bound Rap1, is fused to dTomato (dTom-RglRA); Guanine nucleotide Exchange Factors (GEFs) and GTPase Activating Proteins (GAPs) activate or inactivate Rap1, respectively. Domain organization of full-length Rgl (REM, RasGEF, RA) is shown below. See Fig. 2 for characterization of the biosensor. (B) Surface views of embryos ubiquitously expressing the Rap1 biosensor in control and maternal Rap1-RNAi at the end of cellularization. In control, the biosensor outlines cell membranes; when Rap1 is depleted, cortical signal is reduced and cell outlines are no longer resolved, confirming functional knockdown. Images are autoscaled independently to show the spatial distributions, so intensities are not directly comparable between panels; quantification of biosensor response to altered Rap1 levels and activity is shown in Fig. 2C. n = 10 embryos per condition. Scale bars, 20 µm.

**Figure 1—Supplement 2.**
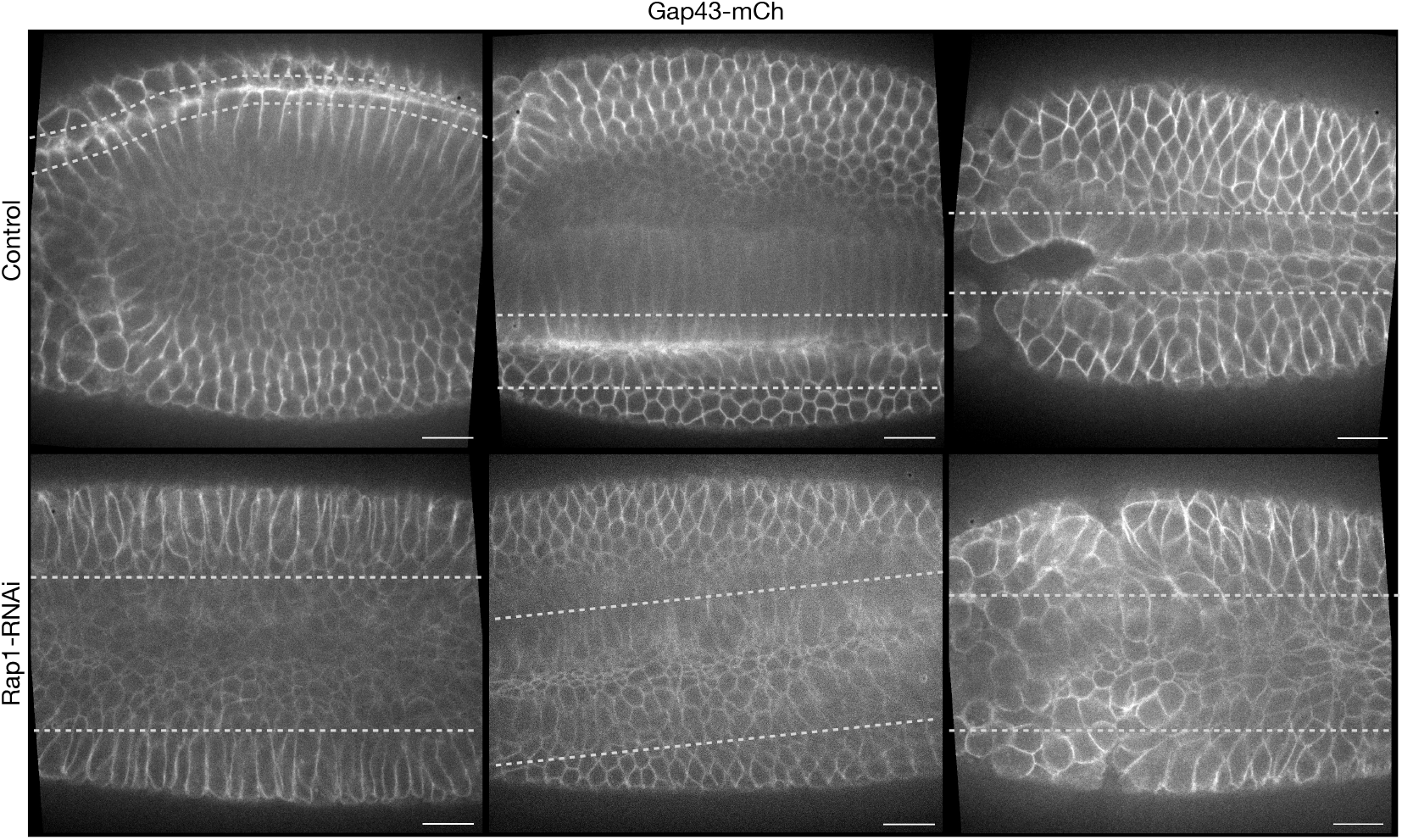
Maternal Rap1-RNAi leads to ventral furrow failure. Ventral views of control (top) and maternal Rap1-RNAi (bottom) embryos expressing Gap43-mCherry (plasma membrane). Each panel is a different embryo. Dashed lines flank the completed furrow or region attempting to make a furrow. n = 5 control and 8 Rap1-RNAi embryos. Scale bar, 20 µm.

**Figure 2—Supplement 1.**
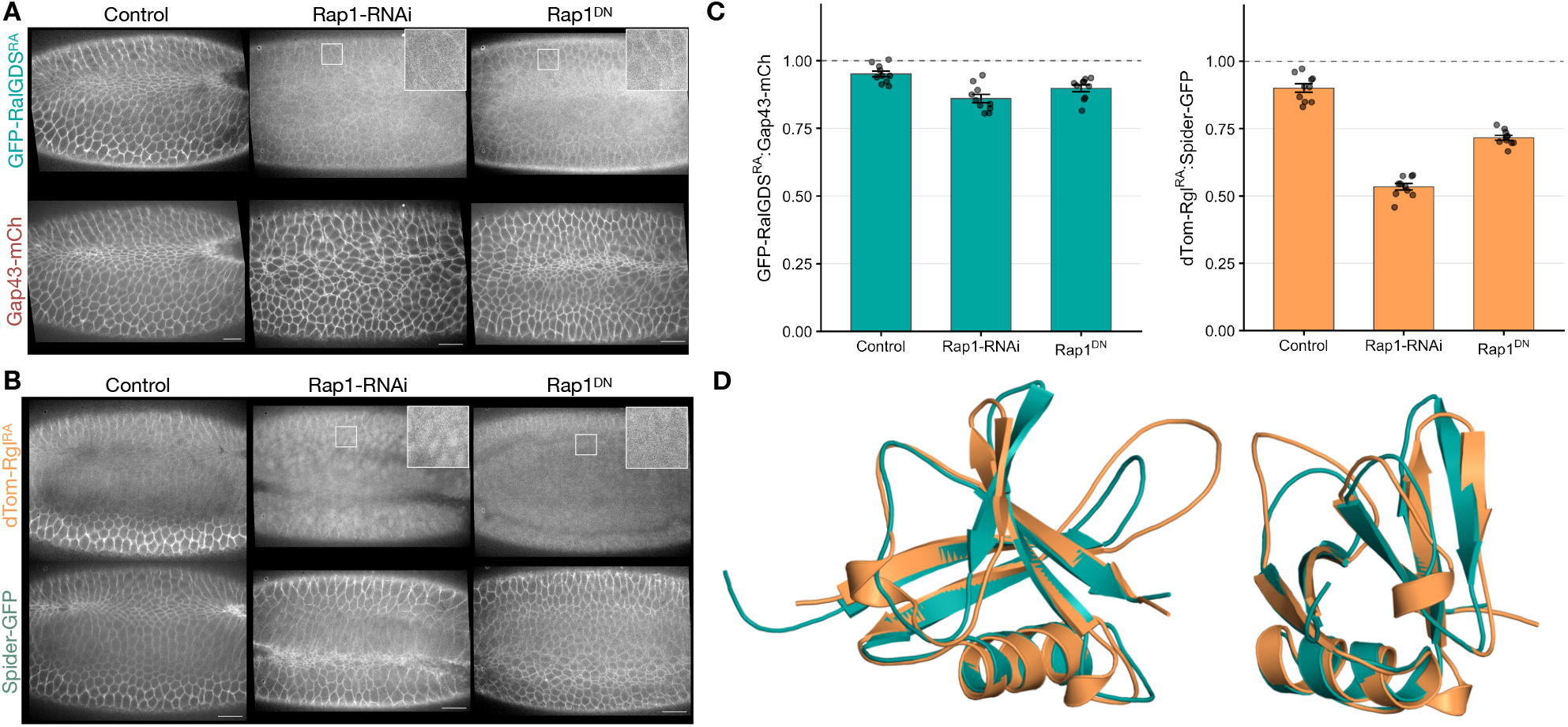
dTom–Rgl^RA^ exhibits improved Rap1-dependent cortical recruitment relative to the RalGDS^RA^-based biosensor in the early *Drosophila* embryo. (A) Ventral views of embryos expressing GFP-RalGDS^RA^ (top) and the membrane marker Gap43-mCherry (bottom) in control embryos and in response to maternal Rap1-RNAi or Rap1^DN^ expression. Insets, cortical GFP-RalGDS^RA^ signal in response to each perturbation. Insets show the boxed regions at 3x magnification. Scale bars, 20 µm. (B) As in A, dTom-Rgl^RA^ (top) with the membrane marker Spider-GFP (bottom). Insets, the stronger loss of cortical dTom-Rgl^RA^ recruitment in response to Rap1 depletion or inhibition. Insets show the boxed regions at 3x magnification. Scale bars, 20 µm. C) Cortex:cytoplasm ratio of each biosensor normalized to that of its co-imaged membrane marker: GFP-RalGDS^RA^ to Gap43-mCherry (teal), dTom-Rgl^RA^ to Spider-GFP (orange). Bars, mean; error bars, SEM; points, individual embryos. n = 10 embryos per condition. (D) Structural comparison of the RA domains of *Drosophila* Rgl (orange; AlphaFold DB model AF-Q9U4U5-F1, residues 858–948) and human RalGDS (teal; solution NMR ensemble PDB 1RAX, residues 788–884). Left, full superposition; right, rotated view. Structures were superposed on Cα atoms with TM-align. The two domains share the ubiquitin-like RA fold (TM-score 0.838; Cα RMSD 1.48 Å over 85 aligned pairs at 42.4% sequence identity), and the fold assignment is robust across all 10 NMR conformers (TM-score 0.817–0.838). Deviation is confined to loops and termini, with 77 of 85 pairs superposing to within 2 Å.

**Figure 3—Supplement 1.**
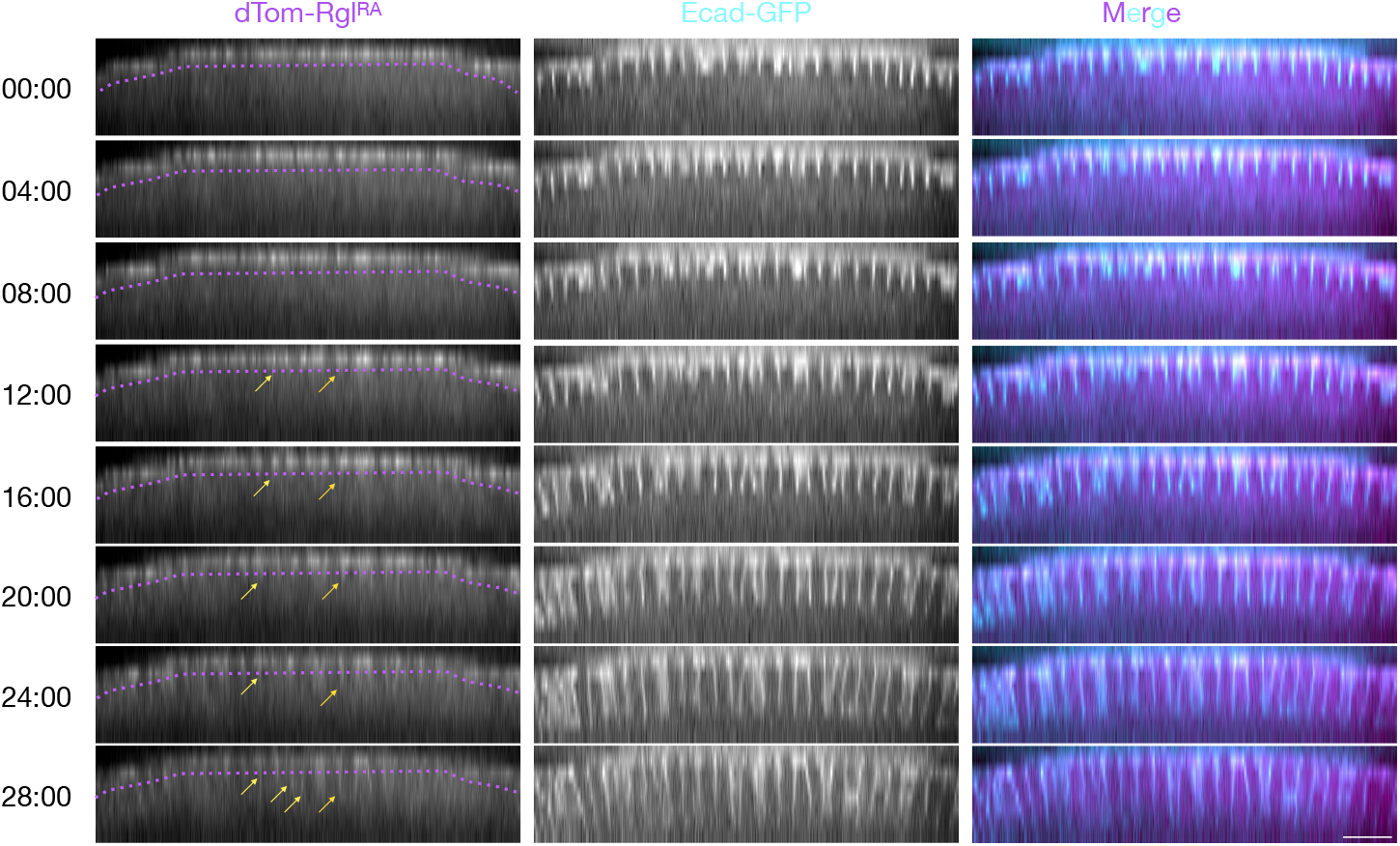
The Rap1 biosensor is enriched apicolaterally during cellularization, with a faint basal pool. XZ views of an embryo co-expressing the Rap1 biosensor (left) and E-cad:GFP (middle); merge at right (biosensor, magenta; E-cadherin, cyan). Apical is up. Dotted magenta line is placed below the apicolateral position to distinguish between the apical and basal Rap1 biosensor signals. Arrows, faint biosensor signal at the basal ends of the ingressing membranes. Time (mm:ss) is relative to the start of acquisition, which began during the slow phase of cellularization. Scale bar, 20 µm; applies to all panels. n = 10 embryos.

**Figure 4—Supplement 1.**
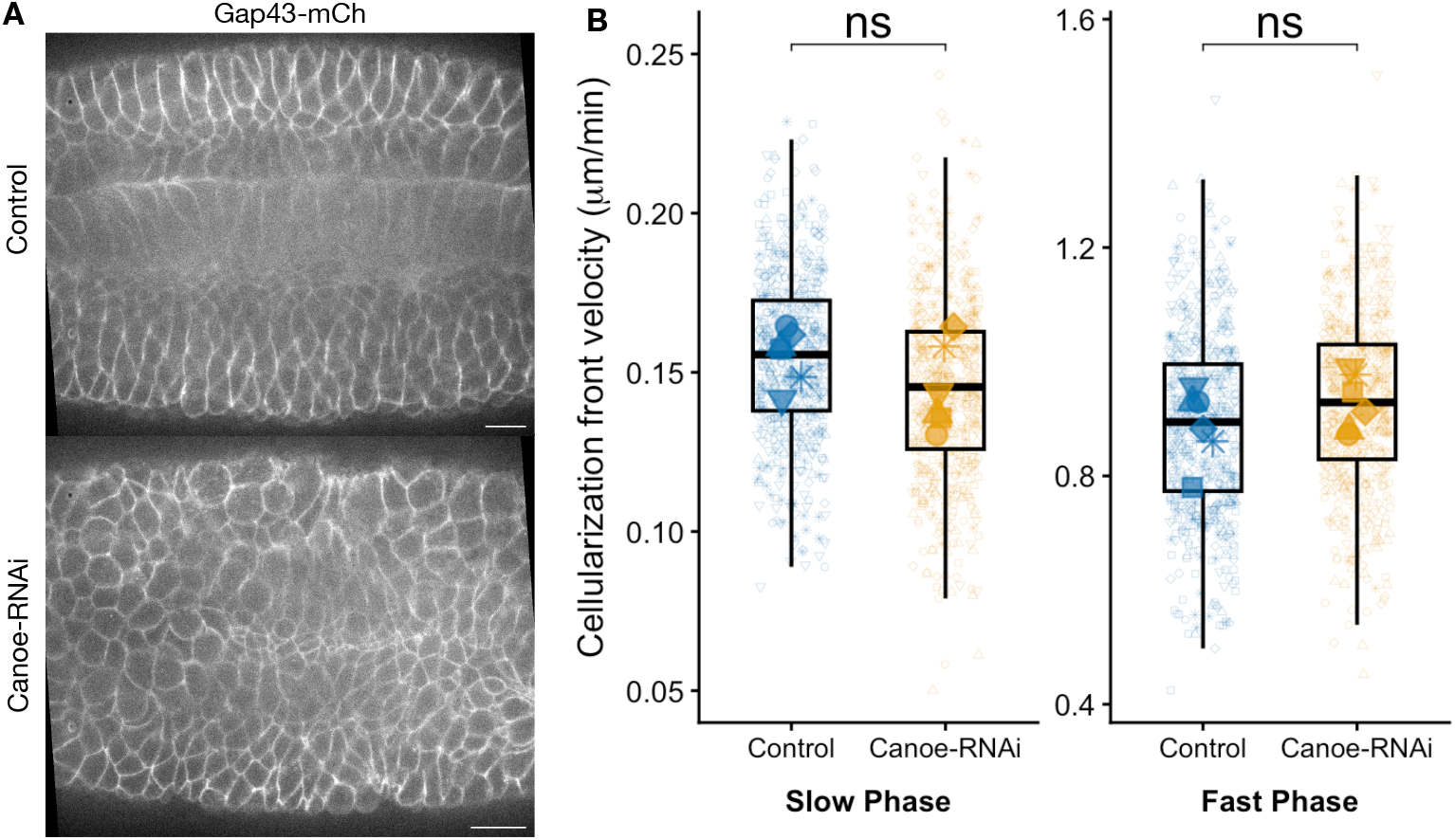
Maternal Canoe depletion disrupts ventral furrow formation without altering the rate of cellularization. (A) Ventral views of embryos expressing Gap43-mCherry as a membrane marker, in control embryos (top) and in response to maternal Canoe depletion (bottom), shown at the initiation of germ-band extension. Scale bars, 20 µm. (B) Cellularization front ingression rate during the slow (left) and fast (right) phases in control (blue) and Canoe-depleted (orange) embryos. Boxes, median and IQR; small symbols, individual front measurements (100 fronts per embryo), with shape denoting the embryo of origin within each condition; large symbols, per-embryo means. Statistics are Welch’s t-tests on per-embryo means. n = 6 embryos per condition. Note the different y-axis scales for the two phases.

**Figure 4—Supplement 2.**
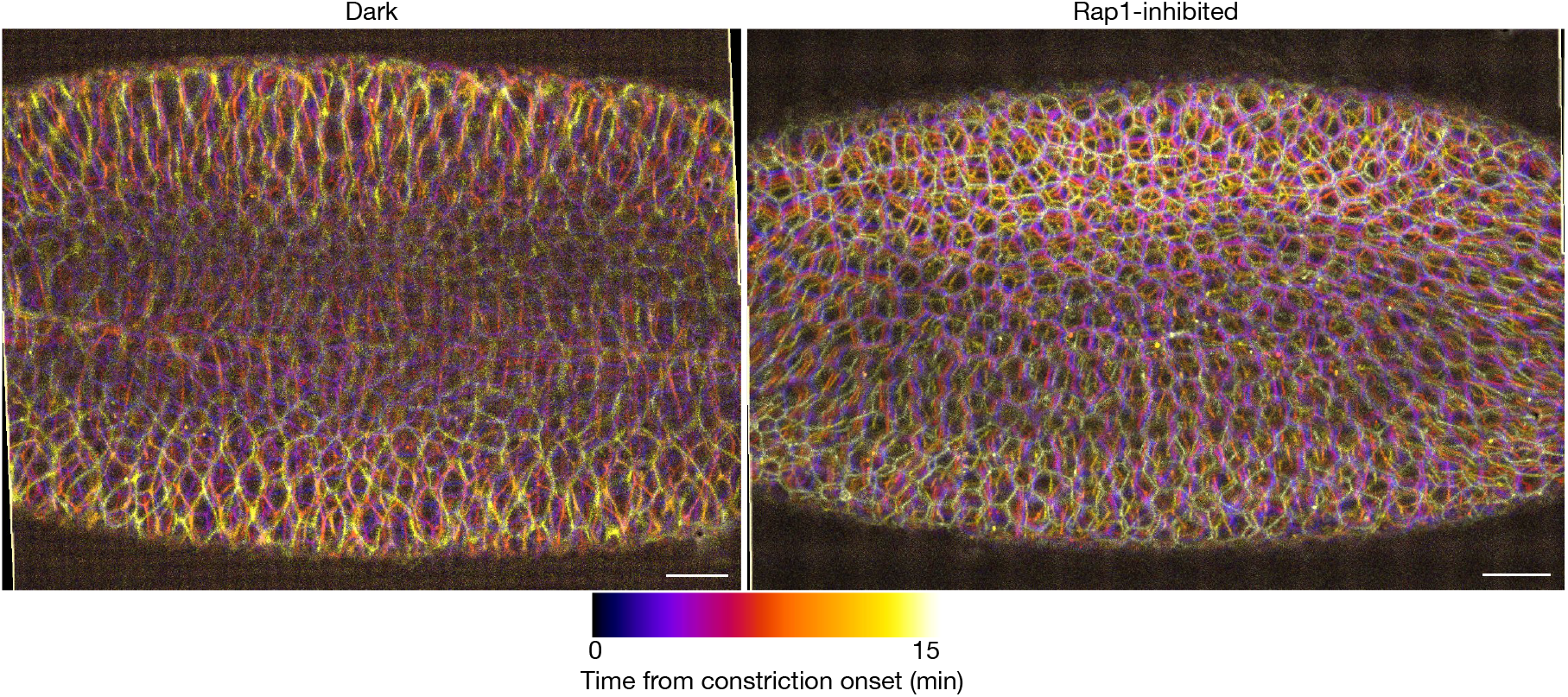
Ventral furrow formation overlaps with germ-band extension in Rap1 inhibited embryos. Temporal color-coded overlays of ventral views of optoRap1^DN^ embryos co-expressing Gap43-mCherry, in dark-control conditions (left) or global Rap1 inhibition from cellularization onset (right). Six frames spanning 15 min, at 3-min intervals, were overlaid with each frame assigned a color according to its time point (Fire LUT; see scale). Both series begin at the onset of apical constriction in that embryo, so the two panels cover the same interval relative to constriction but different absolute developmental times. Structures that do not move over the interval appear white; structures that move appear as separated color fringes. Background was subtracted with a 50-pixel rolling-ball filter. Scale bars, 20 µm. n = 5 embryos per condition.

**Figure 4—Supplement 3.**
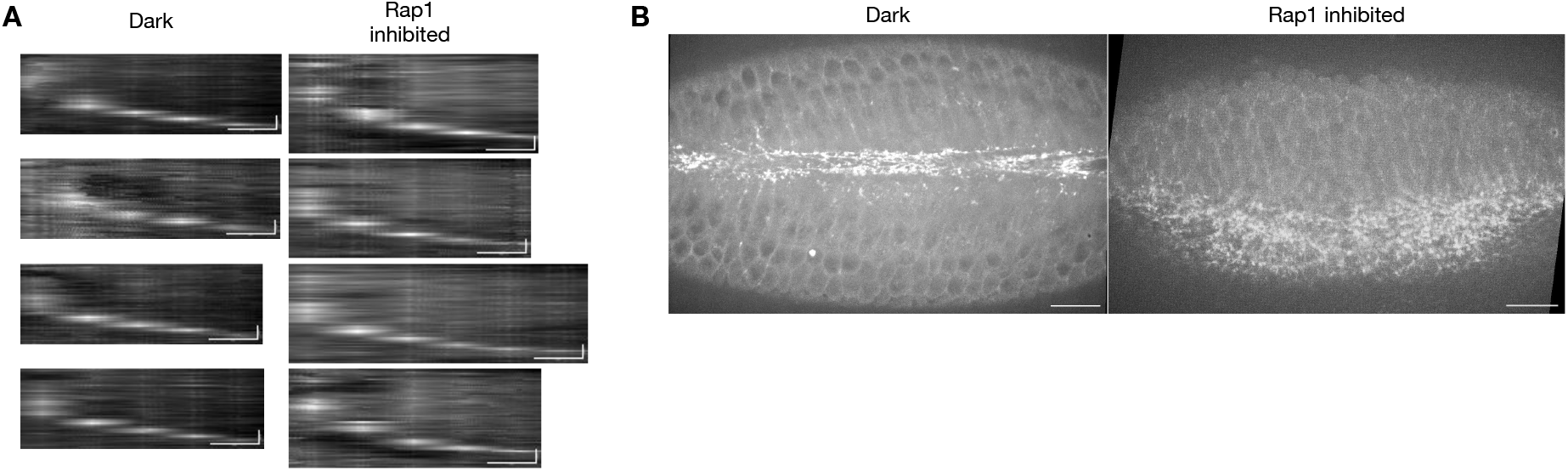
Myosin remains associated with the ingressing front and subsequently accumulates into apical foci in response to Rap1 inhibition. (A) Kymographs of myosin (Sqh-mCherry) at four representative individual cellularization fronts, each from a different embryo, in dark-control (left) and Rap1-inhibited (right) embryos. Each kymograph was generated from a line drawn along the apicobasal axis through a single front, with depth on the horizontal axis and time running downward from t = 0 at the onset of cellularization. Scale bars, 5 µm (horizontal) and 10 min (vertical). n = 5 embryos analyzed per condition. (Note that the Rap1-inhibited kymographs are longer because of slow cellularization.) (B) Maximum-intensity projections of apical sections 5 min after the onset of apical constriction, in dark-control (left) and Rap1-inhibited (right) embryos. Display settings are matched between conditions. Scale bars, 20 µm. n = 5 embryos per condition.

**Figure 5—Supplement 1.**
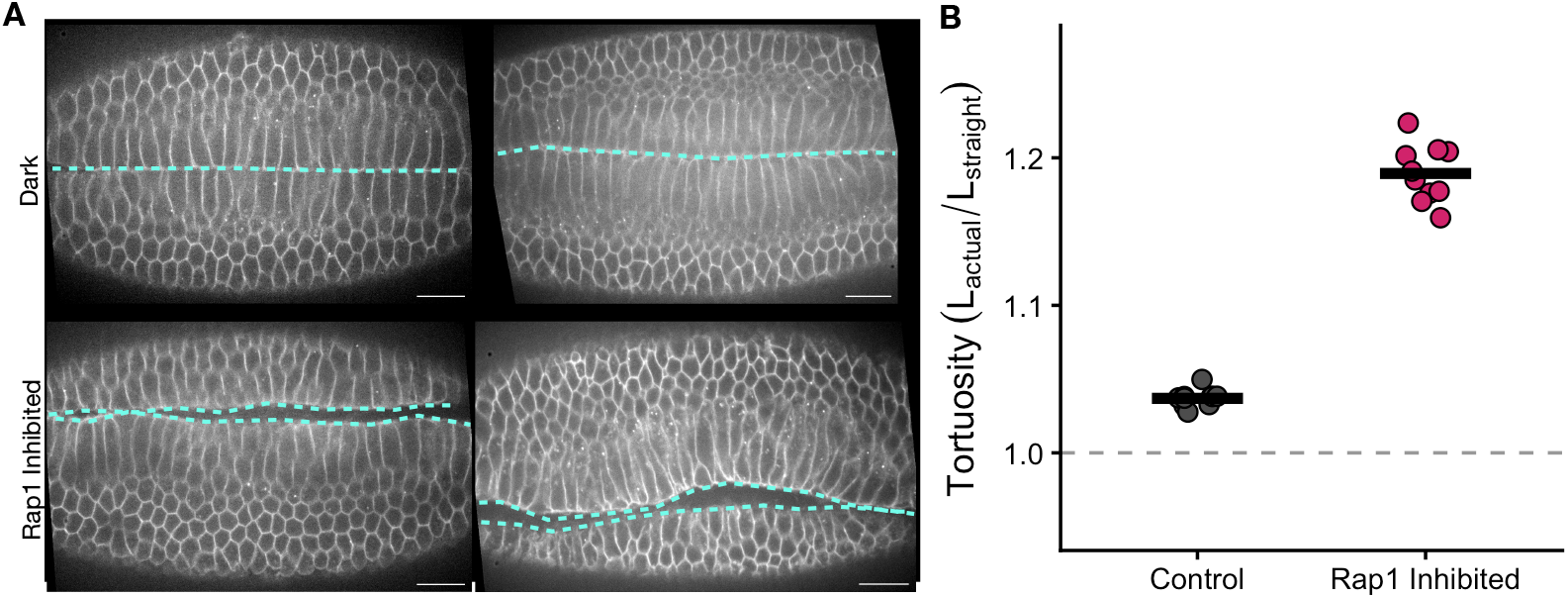
Furrow sealing is slower and follows a more irregular line in response to acute Rap1 inhibition during furrowing. (A) Ventral views of optoRap1^DN^ embryos co-expressing Gap43-mCherry, under dark-control conditions (top) or Rap1 inhibition from VFF onset (bottom); two embryos per condition, at a matched time of 13 min after end of cellularization. By this point, dark controls have sealed along a single line, whereas Rap1-inhibited embryos are still sealing, with the two apposing edges visible. Cyan dashed lines, manually traced seal (dark controls) or apposing furrow edges (Rap1-inhibited). Scale bars, 20 µm. (B) Tortuosity of the traced line, defined as traced path length divided by end-to-end distance, so that a perfectly straight line gives 1.0 (dashed gray line). Each point is one embryo; for Rap1-inhibited embryos, the two apposing edges were traced separately and averaged. Horizontal bars, mean. n = 10 embryos in each condition.

**Figure 6 — Supplement 1.**
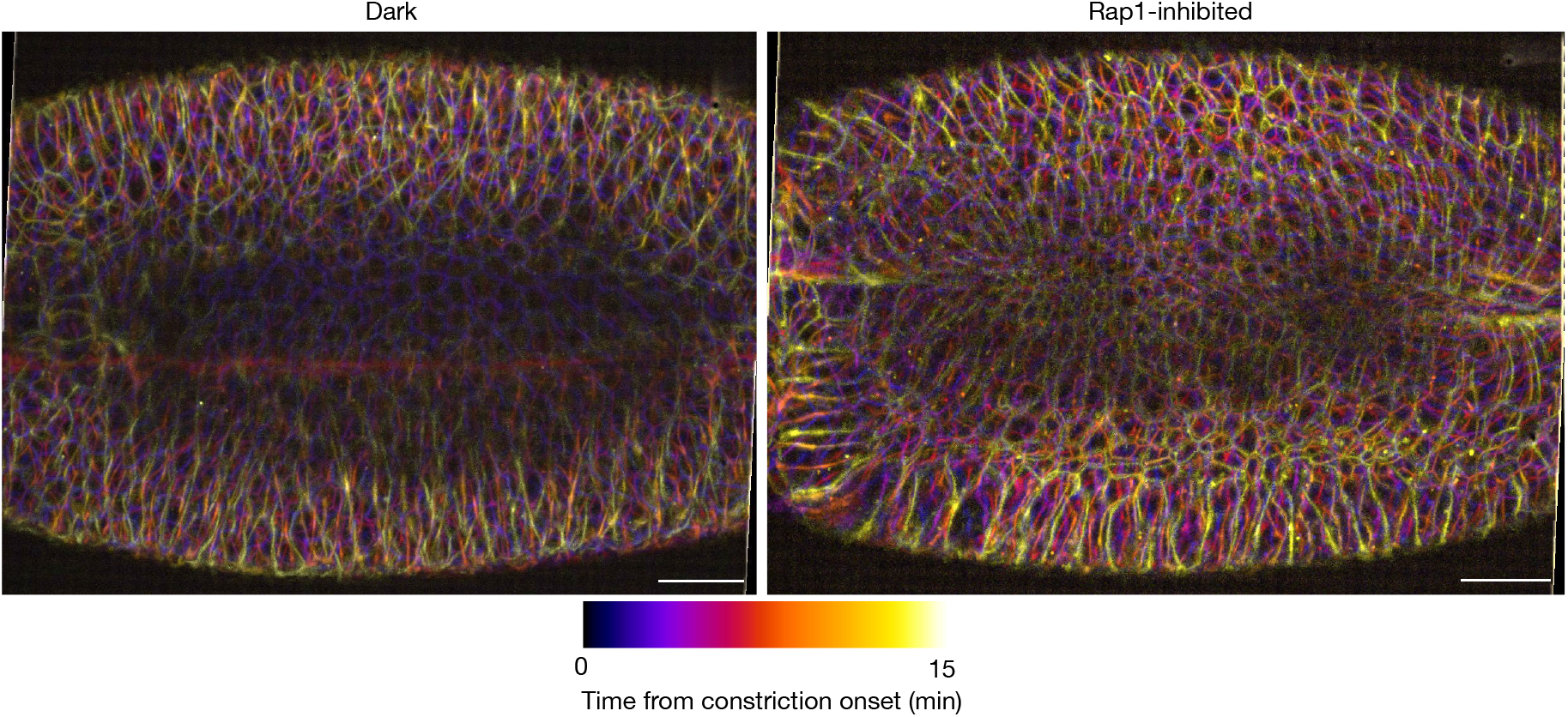
Ventral furrow formation overlaps germ-band extension in response to optogenetic Inhibition of Rap1. Temporal color-coded overlays of ventral views of optoRap1^DN^ embryos co-expressing Gap43-mCherry, under dark-control conditions (left) or global Rap1 inhibition restricted to early cellularization (right). Six frames spanning 15 min, at 3-min intervals, were overlaid with each frame assigned a color according to its timepoint (Fire LUT; see scale). Both series begin at the onset of apical constriction in that embryo, so the two panels cover the same interval relative to constriction but different absolute developmental times. Structures that do not move over the interval appear white, where all timepoints superimpose; structures that move appear as separated color fringes. Background was subtracted with a 50-pixel rolling-ball filter. Scale bars, 20 µm. n = 5 embryos per condition.

**Figure 6—Supplement 2.**
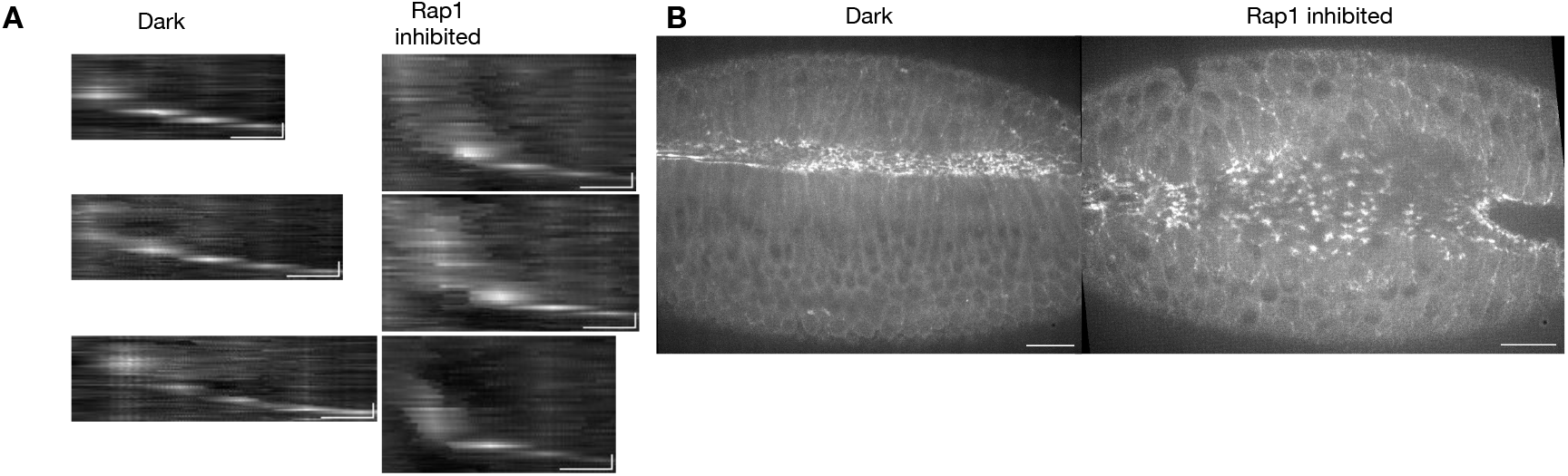
Myosin remains associated with the ingressing front and accumulates into apical foci in response to optogenetic Rap1 inhibition restricted to early cellularization. (A) Kymographs of myosin (Sqh-mCherry) at three representative individual cellularization fronts, each from a different embryo, in dark-control (left) and Rap1-inhibited (right) embryos. Each kymograph was generated from a line drawn along the apicobasal axis through a single front, with depth on the horizontal axis and time running downward from t = 0 at the onset of cellularization. Scale bars, 5 µm (horizontal) and 10 min (vertical). n = 5 embryos analyzed per condition. (Note that the Rap1-inhibited kymographs run longer because of the slow cellularization.) (B) Maximum-intensity projections of apical sections 5 min after the onset of apical constriction, in dark-control (left) and Rap1-inhibited (right) embryos. Display settings are matched between conditions. Scale bars, 20 µm. n = 5 embryos per condition.

**Figure 6—Supplement 3.**
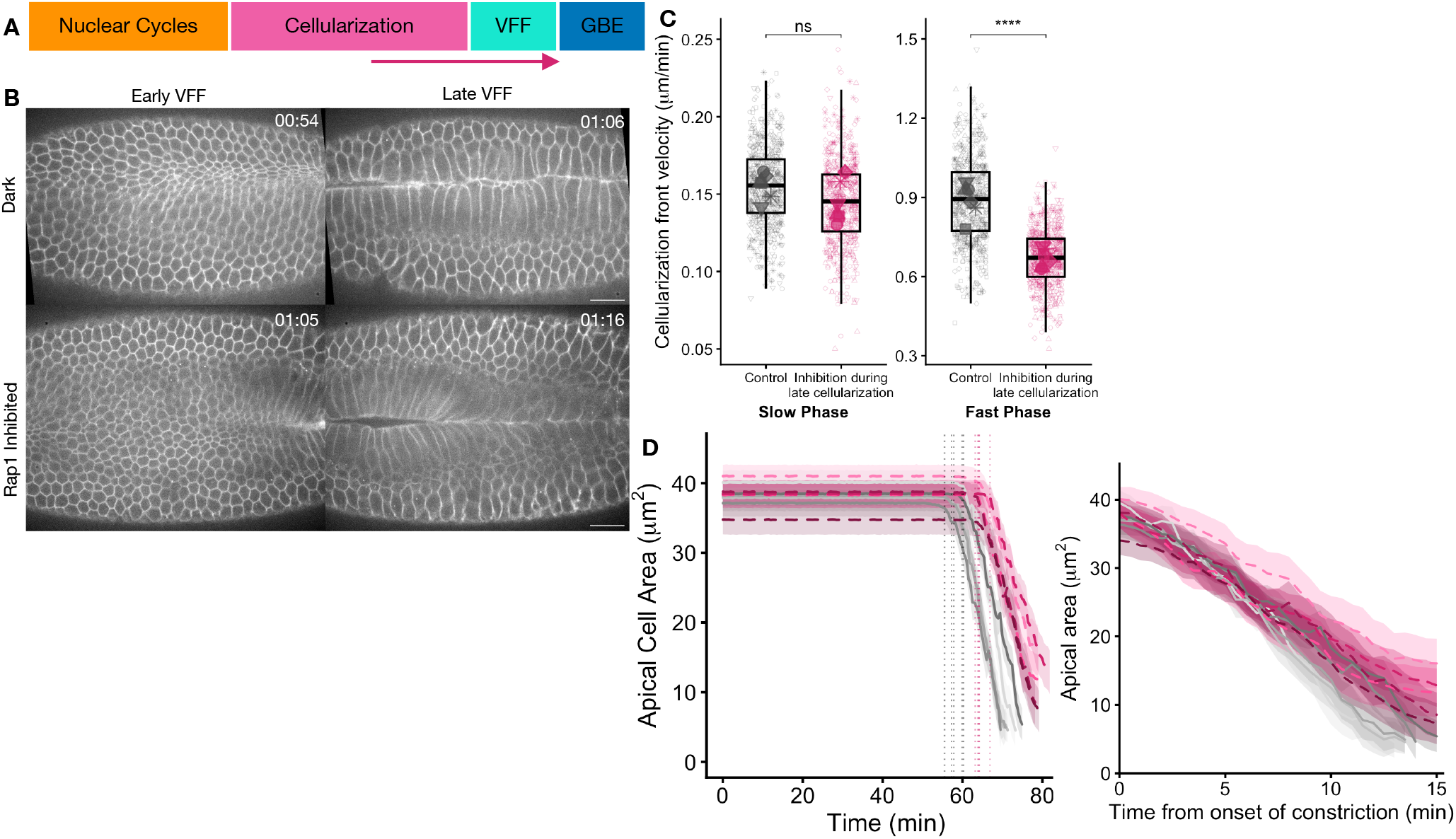
Transient optogenetic Rap1 inhibition during late cellularization does not disrupt VFF. (A) Schematic of early embryonic development marking the illumination window (yellow arrow), which spans the late phase of cellularization after the fronts reach a depth of 10 µm and continues up to VFF. (B) Time-lapse images of optoRap1^DN^ embryos co-expressing Gap43-mCherry, under dark-control conditions (top) or Rap1 inhibition restricted to early cellularization (bottom). Frames are shown at early and late VFF. Time (hh:mm) relative to the end of nuclear cycle 13. Scale bars, 20 µm. (C) Cellularization front velocity during the slow and fast phases in dark controls (gray) and embryos inhibited throughout late cellularization and VFF (magenta). Boxes, median and IQR; small symbols, individual front measurements, with shape denoting the embryo of origin within each condition; large symbols, per-embryo means. Welch’s t-tests on per-embryo means. (D) Apical cell area over time in dark-control (gray, solid) and Rap1-inhibited (magenta, dashed) embryos. Top, traces aligned to the onset of cellularization; dotted vertical lines mark the onset of apical constriction in each embryo, colored by condition and defined as a 5% decrease in apical area. Bottom, the same traces realigned to constriction onset. Each line is the mean apical area of one embryo; shaded bands, SD across cells within that embryo. n = 5 embryos per condition.

**Figure 7—Supplement 1.**
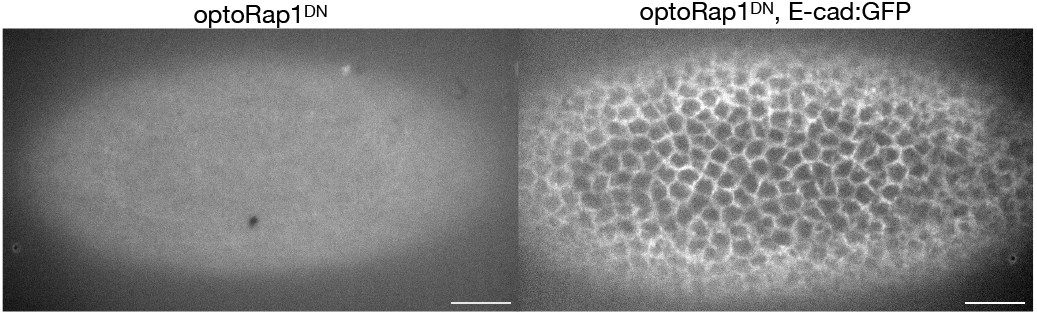
The GFP signal from optoRap1^DN^ is negligible relative to E-cad:GFP. Sum projections of apical z-slices from embryos expressing optoRap1^DN^ alone (left) or optoRap1^DN^ together with E-cad:GFP (right), during cellularization. Both were imaged with identical 488 nm excitation and detection settings and are displayed on the same intensity scale. 3 slices spanning 10 µm from the apical surface were summed. Scale bars, 20 µm. n = 5 embryos per genotype.

**Supplementary File 1.**
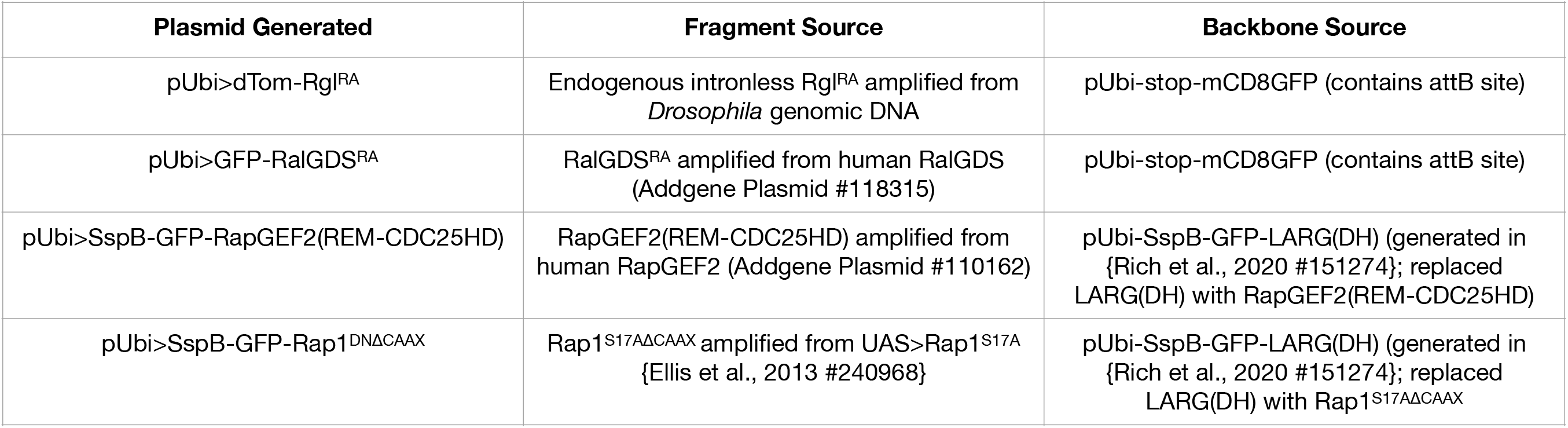

**Supplementary File 2.**
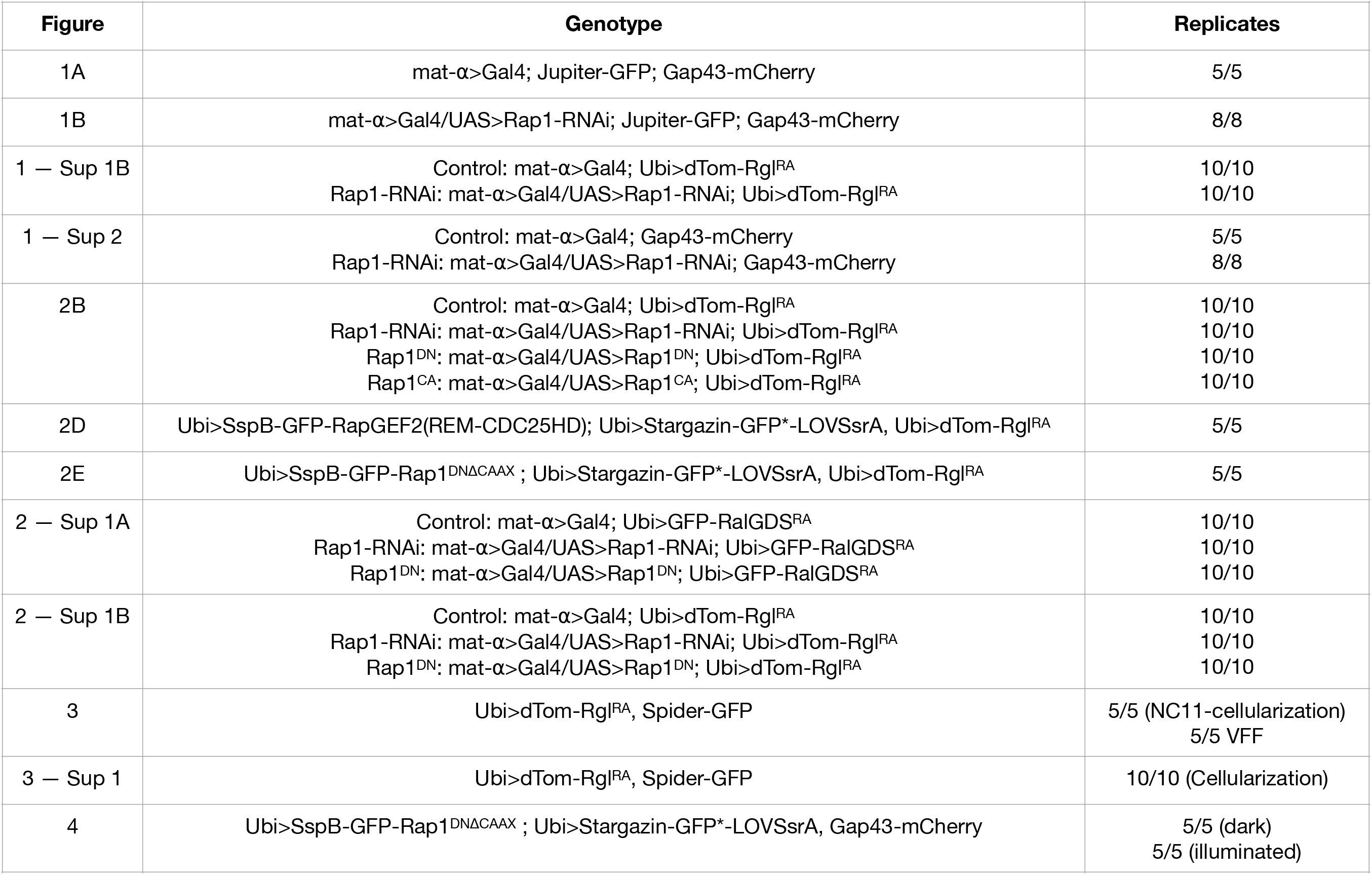
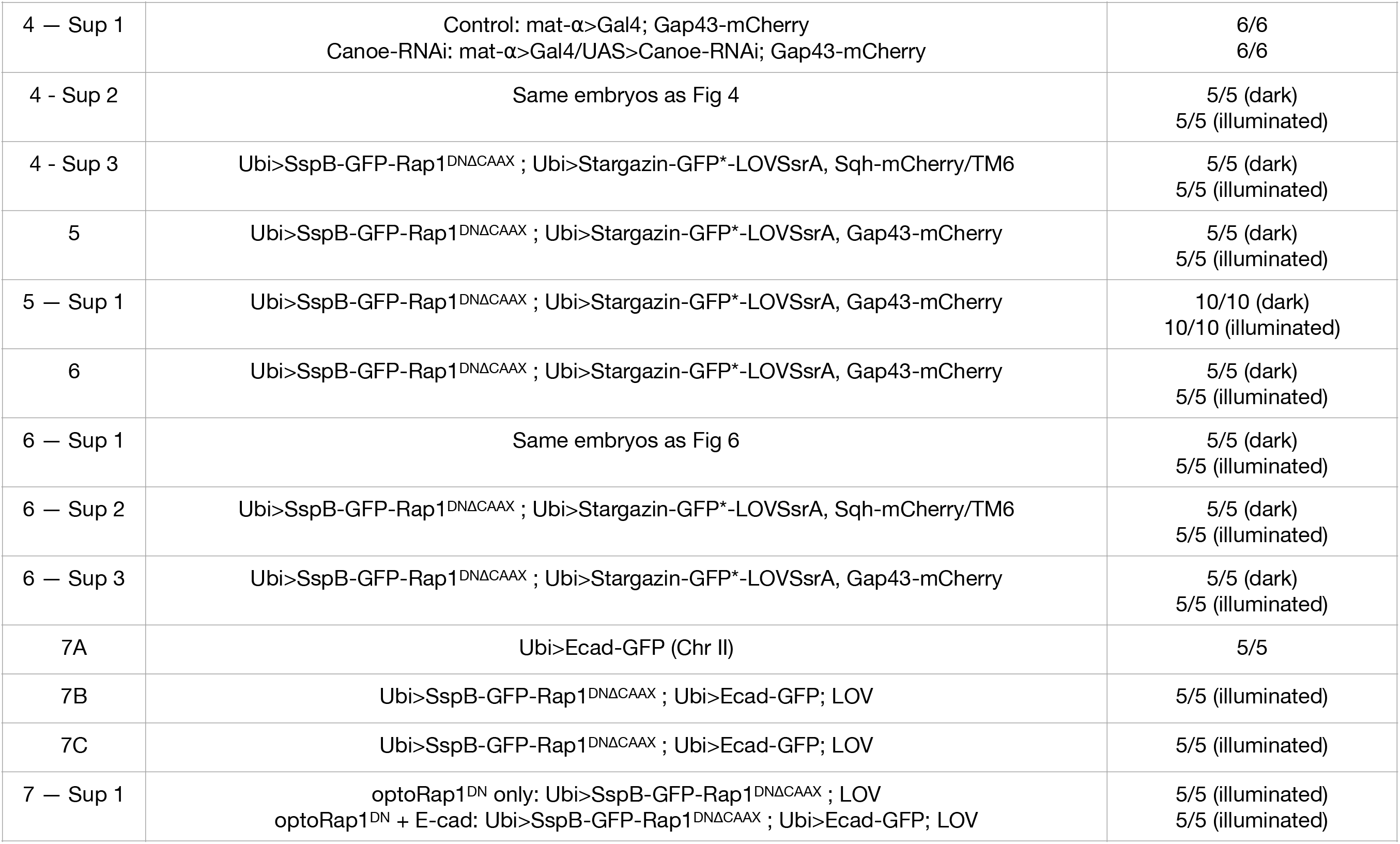

**Supplementary File 3.**
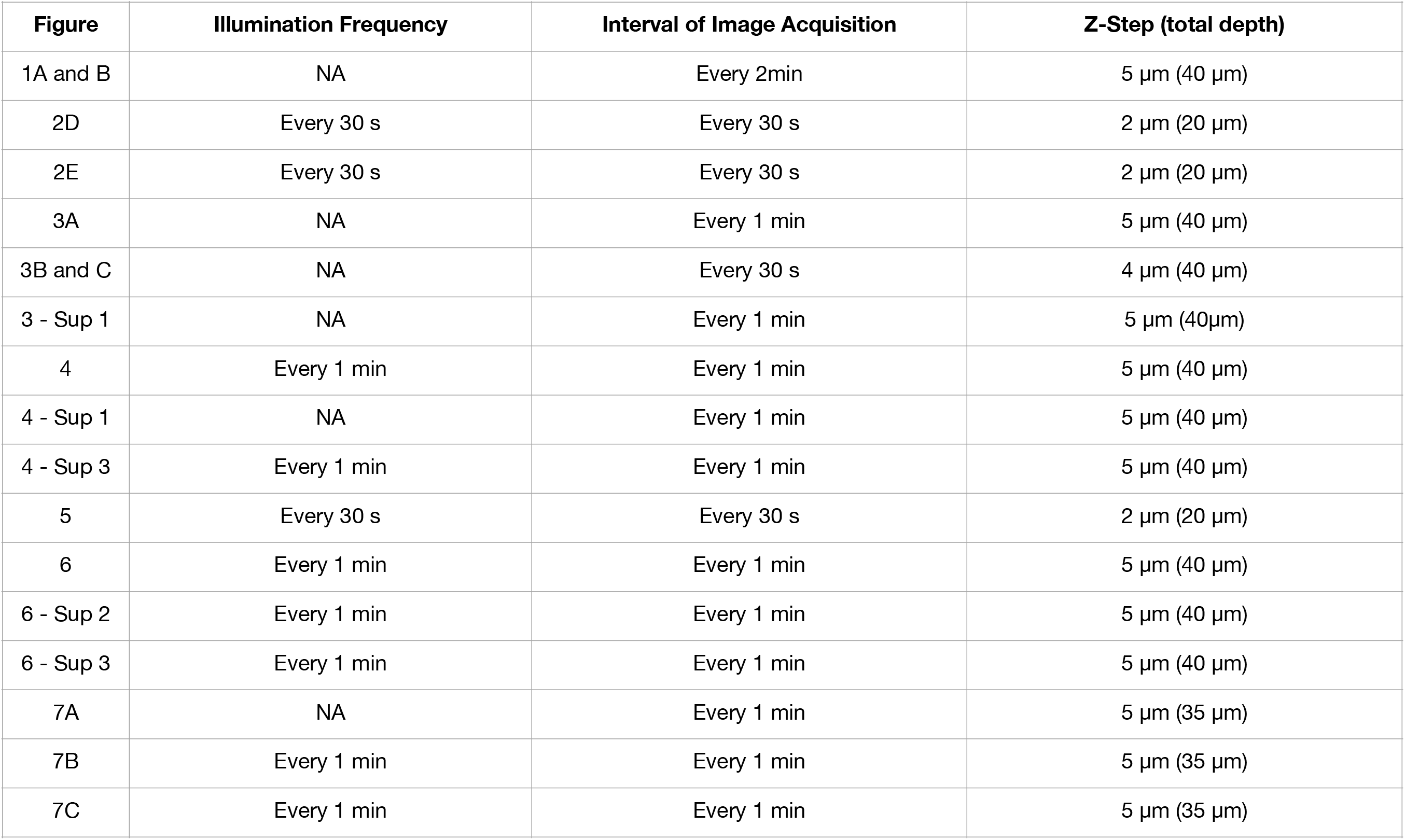
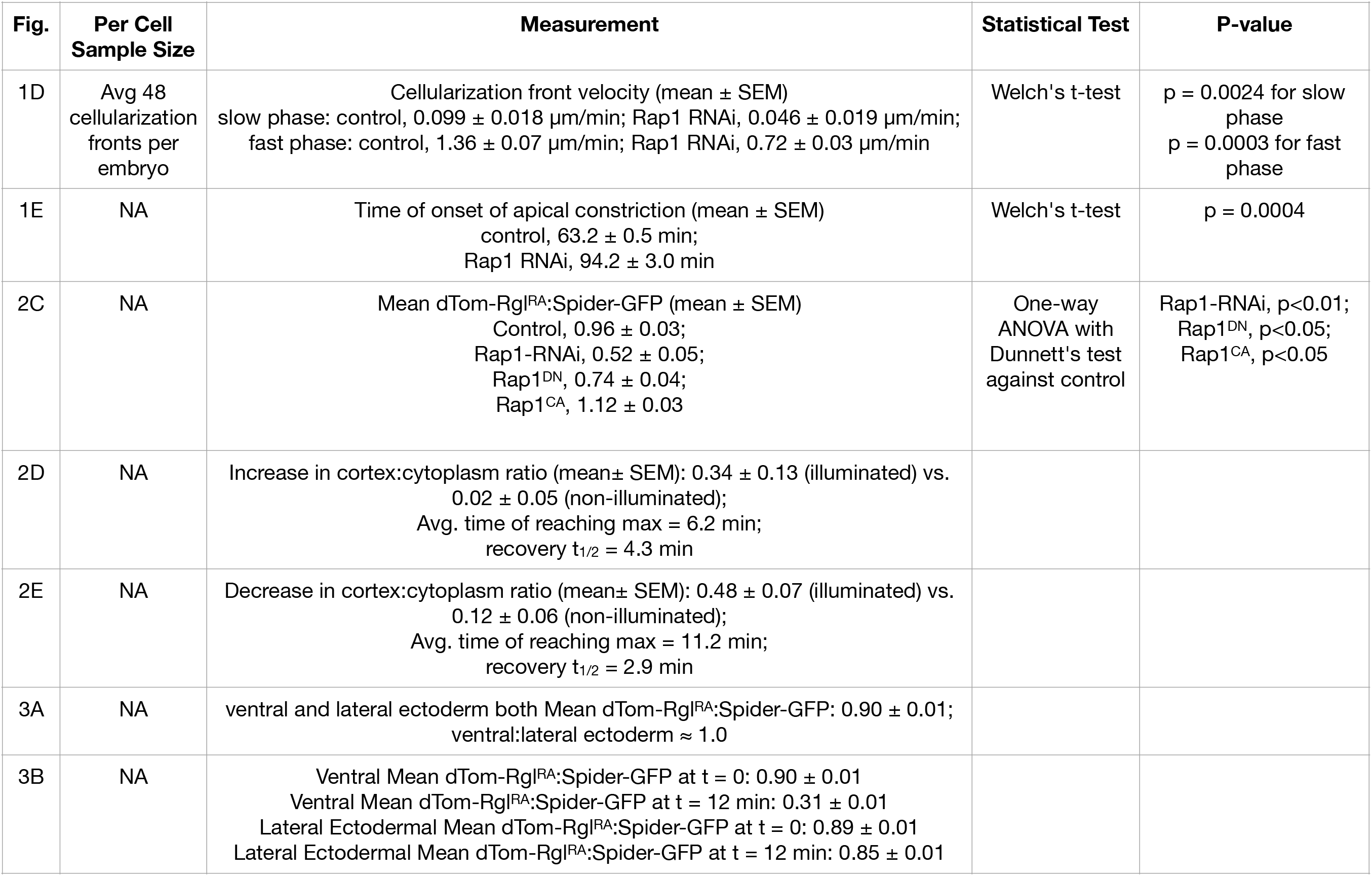
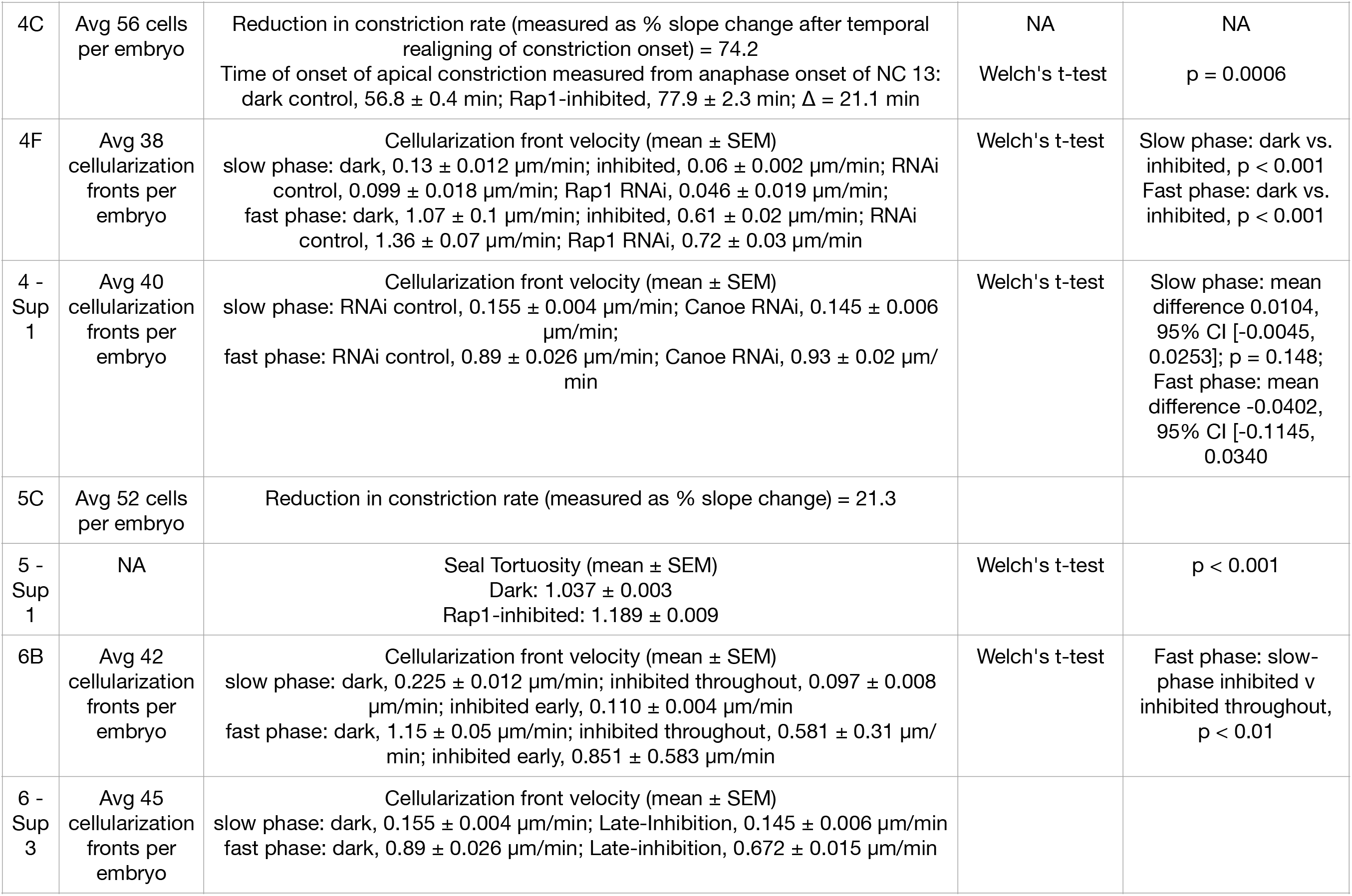
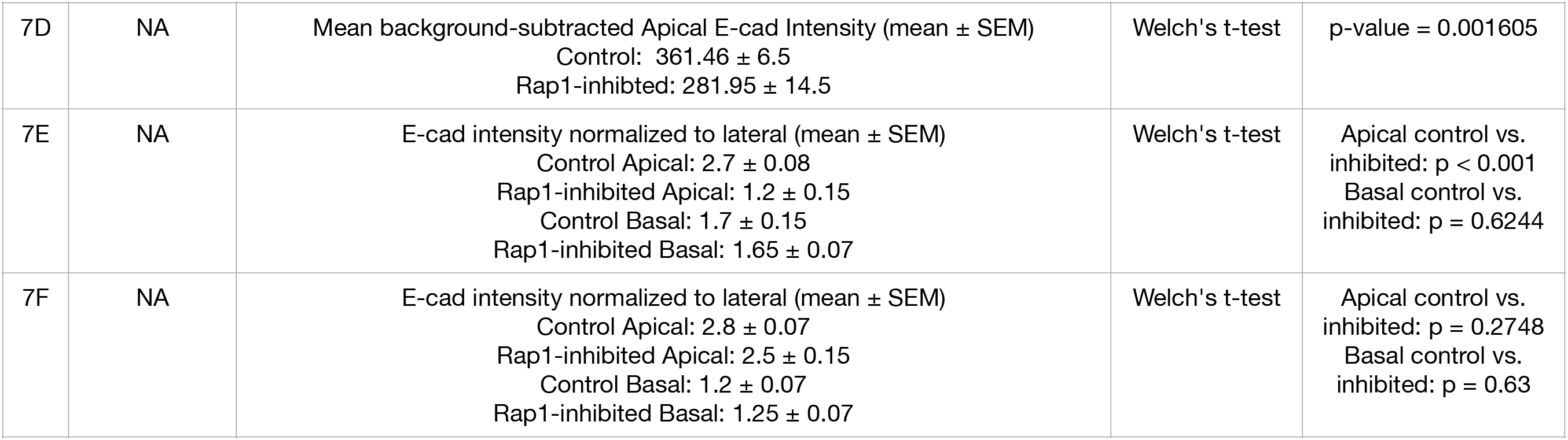

**Video S1:** Ventrolateral view of a control embryo (driver and markers without an RNAi transgene; full genotype in Supplementary File 2) expressing Jupiter-GFP (microtubules, cyan) and Gap43-mCherry (plasma membrane, magenta). Single plane 15 µm below the embryo surface. Anterior is to the left. Images were acquired every 2 min from anaphase onset of nuclear cycle 13 (nc13; t = 0) until the appearance of the anterior mitotic domain MD4, visible in the lower right of the field. Time is displayed as hh:mm relative to nc13 anaphase onset.

**Video S2:** Ventrolateral view of a Rap1-depleted embryo (driver and markers with Rap1-RNAi transgene; full genotype in Supplementary File 2) expressing Jupiter-GFP (microtubules, cyan) and Gap43-mCherry (plasma membrane, magenta). Single plane 10 µm below the embryo surface. Anterior is to the left. Images were acquired every 2 min from anaphase onset of nuclear cycle 13 (nc13; t = 0). Time is displayed as hh:mm relative to nc13 anaphase onset.

**Video S3:** Single confocal plane 10 µm below the embryo surface in an embryo expressing Ubi>dTom-Rgl^RA^. Ventral view; anterior to the left. Images were acquired every 1 min from nuclear cycle 11 (nc11; t = 0) through the end of nc13; total elapsed time 51 min, displayed as hh:mm.

**Video S4:** Single confocal plane 10 µm below the embryo surface in an embryo expressing Ubi>dTom-Rgl^RA^. Ventral view; anterior to the left. Images were acquired every 1 min from nuclear cycle 13 (nc13; t = 0) through the end of cellularization; total elapsed time 1 h, displayed as hh:mm.

**Video S5.** dTom-Rgl^RA^ during ventral furrow formation. Single confocal plane 12 µm below the embryo surface (left) and YZ orthogonal reslice of the same embryo (right), expressing Ubi>dTom-Rgl^RA^. In the *en face* view, anterior is to the left; in the YZ view, lateral regions above and below and apical is to left, with the ventral midline centered. Images were acquired every 30 s from the end of cellularization (t = 0) through the completion of furrowing; total elapsed time 15 min, displayed as hh:mm.

**Video S6. Membrane dynamics from cellularization onset through ventral furrow formation in a dark control embryo.** Embryo expressing optoRap1^DN^; Gap43-mCherry without any optogenetic perturbation. Single confocal plane 20 µm below the embryo surface (left) and YZ orthogonal reslice of the same embryo (right). In the *en face* view, anterior is to the left; in the YZ view, apical is to the left, with the ventral midline centered and lateral regions above and below. Images were acquired every 1 min from the onset of cellularization (t = 0) through the completion of furrowing; total elapsed time 65 min, displayed as hh:mm.

**Video S7. Membrane dynamics in a Rap1-inhibited embryo perturbed throughout cellularization and VFF.** Embryo expressing optoRap1^DN^; Gap43-mCherry with optogenetic perturbation throughout timelapse acquisition. Single confocal plane 35 µm below the embryo surface (left) and YZ orthogonal reslice of the same embryo (right). In the *en face* view, anterior is to the left; in the YZ view, apical is to the left, with the ventral midline centered and lateral regions above and below. Images were acquired every 1 min from the onset of cellularization (t = 0) through the completion of furrowing; total elapsed time 72 min, displayed as hh:mm.

**Video S8. Membrane dynamics during ventral furrow formation in a dark control embryo.** Embryo expressing optoRap1^DN^; Gap43-mCherry with no optogenetic perturbation. Single confocal plane 14 µm below the embryo surface. Anterior is to the left. Images were acquired every 30 seconds from the end of cellularization (t = 0) through the completion of furrowing; total elapsed time 17 min, displayed as mm:ss.

**Video S9. Membrane dynamics in a Rap1-inhibited embryo perturbed throughout VFF.** Embryo expressing optoRap1^DN^; Gap43-mCherry with optogenetic perturbation throughout timelapse acquisition. Single confocal plane 14 µm below the embryo surface (left) and YZ orthogonal reslice of the same embryo (right). In the *en face* view, anterior is to the left; in the YZ view, apical is to the left, with the ventral midline centered and lateral regions above and below. Images were acquired every 30 seconds from the end of cellularization (t = 0) through the completion of furrowing; total elapsed time 17 min, displayed as mm:ss.

**Video S10. Membrane dynamics in a Rap1-inhibited embryo perturbed only during early cellularization and imaged up to VFF.** Embryo expressing optoRap1^DN^; Gap43-mCherry with optogenetic perturbation only during early cellularization (indicated with overlay). Single confocal plane 40 µm below the embryo surface (left) and YZ orthogonal reslice of the same embryo (right). In the *en face* view, anterior is to the left; in the YZ view, apical is to the left, with the ventral midline centered and lateral regions above and below. Images were acquired every 1 min from the onset of cellularization (t = 0) through the completion of furrowing; total elapsed time 79 min, displayed as hh:mm.

**Video S11. Myosin localization from cellularization onset through ventral furrow formation in a dark-control embryo.** Embryo expressing optoRap1DN and Sqh-mCherry, imaged with 561 nm illumination only, which does not activate the LOV domain. Single confocal plane 35 µm below the embryo surface (left). Anterior is to the left. Images were acquired every 1 min from the onset of cellularization (t = 0) through the completion of furrowing; total elapsed time 66 min, displayed as hh:mm.

**Video S12. Myosin localization from cellularization onset through ventral furrow formation in a Rap1-inhibited embryo perturbed only during early cellularization.** Embryo expressing optoRap1^DN^; Sqh-mCherry with optogenetic perturbation only during early cellularization (indicated with overlay). Single confocal plane 40 µm (basal view) below the embryo surface; anterior is to the left. Images were acquired every 1 min from the onset of cellularization (t = 0) through the completion of furrowing; total elapsed time 129 min, displayed as hh:mm.

**Video S13. Myosin localization from cellularization onset through ventral furrow formation in a Rap1-inhibited embryo perturbed only during early cellularization.** Same embryo as Video 12 expressing optoRap1^DN^; Sqh-mCherry with optogenetic perturbation only during early cellularization (indicated with overlay). Sum of single confocal plane with total depth of 15 µm below the embryo surface; anterior is to the left. Images were acquired every 1 min from the onset of cellularization (t = 0) through the completion of furrowing; total elapsed time 129 min, displayed as hh:mm.

